# *Streptomyces* induces unisexual reproduction of a major human fungal pathogen

**DOI:** 10.64898/2026.10.05.756715

**Authors:** Ye-Eun Son, Xuefei Chen, Gerard D. Wright, Sheng Sun, Joseph Heitman

## Abstract

*Cryptococcus*, a human pathogenic yeast, can undergo unisexual reproduction in the absence of compatible mating partners, yet this developmental program remains poorly understood. Here, we report that tunicamycin, a natural product produced by *Streptomyces* species, induces a complete unisexual reproductive cycle in different *Cryptococcus* species. Bacterial confrontation or purified tunicamycin trigger hyphal differentiation, meiotic recombination, and basidiospore formation, as confirmed by whole-genome sequencing and SNP analyses. We further demonstrate that tunicamycin-induced unisexual development requires canonical sexual regulators and meiotic machinery. Transcriptomic profiling reveals staged developmental reprogramming characterized by early induction of stress- and cell cycle–associated genes, followed by enrichment of genes in the filamentation-related pathways. This capacity varies among natural isolates of *C. neoformans* and was conserved in *Cryptococcus gattii*. Together, our findings demonstrate that a bacterial metabolite can activate meiotically competent unisexual reproduction, highlighting interkingdom chemical signaling as a potential driver of fungal evolution in natural environments.

## Introduction

“In nature we never see anything isolated, but everything in connection with something else which is before it, beside it, under it and over it.” (Johann Wolfgang von Goethe)^1^. At microscopic scales, no microbe exists in isolation; every organism, including bacteria, archaea, fungi, and viruses, is embedded in a vast and dynamic web of interactions shaped by both cooperation and competition, influencing each other’s survival and evolution. Inter-kingdom interactions within these communities can profoundly reshape cellular morphology, developmental programs, and survival strategies^2^.

Previous studies have revealed numerous instances in which crosstalk between bacteria and fungi profoundly influences fungal cell biology and pathogenesis on either side of the inter-species interaction^3–7^. For example, interactions between the human fungal pathogen *Candida albicans* and the Gram-positive bacterium *Enterococcus faecalis* synergistically enhance host cell damage, in part through direct physical association that promotes bacterial accumulation on host cell surfaces^7^. Similarly, the bacterium *Acinetobacter baumannii* induces capsule and biofilm formation in the human pathogen *Cryptococcus neoformans* during cocultivation^3^ and direct cell-to-cell contact between *Pseudomonas aeruginosa* and *C. neoformans* stimulates production of pyocyanin and other inhibitory factors that suppress cryptococcal growth^4^. Bacteria can also influence the sexual behavior of eukaryotes. For example, the marine bacterium *Vibrio fischeri* induces sexual reproduction in the choanoflagellate *Salpingoeca rosetta* through secretion of a chondroitinase^8^. Additionally, *C. elegans* can switch from selfing to mating when pathogenic bacteria are present^9^.

Gram-positive filamentous bacteria of the genus *Streptomyces* are among the most abundant and metabolically versatile microorganisms in soil ecosystems, where they play central roles in nutrient cycling and microbial interactions^10,11^. Members of this genus are prolific producers of bioactive natural products and are estimated to account for approximately two-thirds of all naturally derived microbial agents, including streptomycin, rapamycin, and chloramphenicol, which can act as chemical signals that modulate the physiology, development, and ecological interactions of neighboring microorganisms^12–14^. Together, these properties position *Streptomyces* species as key drivers of interkingdom communication within complex microbial communities.

The *Cryptococcus* species complex comprises basidiomycetous yeasts that cause life-threatening cryptococcal meningoencephalitis, particularly in immunocompromised individuals, such as those living with HIV/AIDS, with reported mortality rates approaching 70%^15–18^. These pathogens are widely distributed in the environment, most commonly in soil enriched with bird guano or decaying wood^19^. Titan cells, a unique morphological form of *Cryptococcus*, are highly resistant to phagocytosis and promote establishment of pulmonary infection and disease progression^16,20^ and sexual reproduction produces sexual basidiospores that function as infectious propagules, resist host oxidative stress, and evade macrophage-mediated clearance^16^. Thus, understanding the environmental cues and regulatory mechanisms that drive these developmental transitions is important not only for fungal biology, but also for understanding cryptococcal pathogenesis.

*Cryptococcus* species exhibit a bipolar mating system where the mating type is determined by a single mating type locus (*MAT*), with isolates belonging to either the *MAT*α or *MAT***a** mating type^21,22^, and employ diverse reproductive strategies, including α × **a** sexual reproduction and pseudosexual reproduction involving cells of opposite mating types, as well as unisexual reproduction involving cells of a single mating type^17,18,23–34^. One enduring conundrum in *Cryptococcus* biology is that although population genomic studies provide strong evidence for ongoing sexual reproduction, isolates collected from most geographic locations are overwhelmingly biased toward the *MAT*α mating type^35–38^. While unisexual reproduction in *Cryptococcus* has thus far been observed primarily under laboratory conditions and is associated with altered G protein signaling and regulation, accumulating evidence suggests that naturally occurring polymorphisms can promote unisexual reproduction, raising the possibility that this reproductive mode occurs more readily and frequently in nature than is currently appreciated^32,37,39^.

Here, we report that tunicamycin produced by two *Streptomyces* isolates closely related to *S. cyanofuscatus* can induce non-yeast cellular development programs, including titan cell formation and unisexual reproduction, in *Cryptococcus* species. This study provides the first demonstration that a bacterially-derived natural product can robustly modulate fungal reproductive strategies, with important implications for understanding the evolution of virulence-associated traits in medically important fungal pathogens.

## Results

### Identification of two *Streptomyces* strains that induce diverse non-yeast development in *C. neoformans*

To explore potential interactions between *C. neoformans* and *Streptomyces* isolates, we first assessed the ability of 39 *Streptomyces* isolates from our in-house collection^40^ to grow on the rich medium YPD, as well as on V8 (pH=5 or pH=7) and MS, two media that support sexual development of *Cryptococcus*. We found that 14 *Streptomyces* isolates failed to grow on MS medium; among these, eight also failed to grow on V8, and three failed to grow on YPD (**Table S1**). Given the higher compatibility on V8 medium, we first used V8 medium to assess how *Streptomyces* isolates might influence cellular development in *C. neoformans*. For each *Streptomyces* isolate, cells from overnight cultures were spotted at the center of a V8 agar plate, with *C. neoformans* wild-type strains H99α, KN99**a**, and KN99α spotted in a circular pattern surrounding the bacterial spot (**Figure 1A**). The co-cultures were incubated at room temperature in the dark, and fungal morphological changes were monitored daily for four weeks. This screening identified two *Streptomyces* isolates, WAC1490 and WAC1565, that consistently induced robust non-yeast cellular development in *C. neoformans* (**Figure 1B**). While solo cultures of *C. neoformans* H99α, KN99**a**, and KN99α only grew as yeast cells on the V8 medium without bacterial cells, they consistently grew as non-yeast forms in the presence of WAC1490 or WAC1565. These forms included enlarged cells, hyphal development, and occasional sporulation (**Figures 1A and 1C**), resembling morphogenesis observed during sexual reproduction in *C. neoformans*.

**Figure 1.**
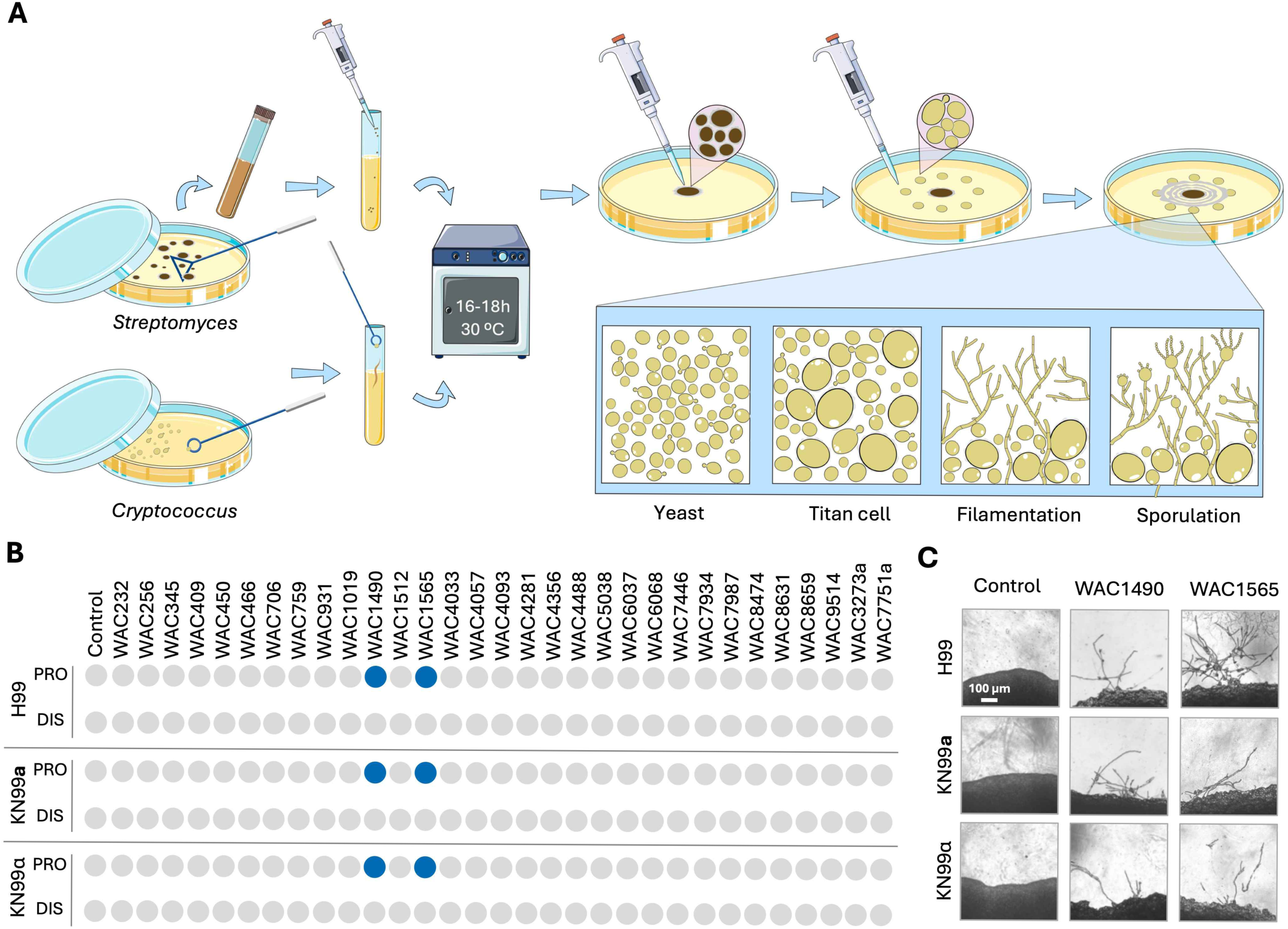
Identification of two *Streptomyces* isolates that induce diverse non-yeast developments in *C. neoformans*. **(A)** Schematic of the experimental design to assess trans-kingdom microbial interactions between *Streptomyces* and *Cryptococcus* species. Both organisms were pre-cultured in liquid YPD medium at 30°C for 16–18 h. A 5 μL aliquot of *Streptomyces* was spotted at the center of the confrontation plate, and 5 μL of *Cryptococcus* cells were spotted at a 1.3 cm distance away from the *Streptomyces* spot. Plates were incubated at room temperature in the dark and observed daily by light microscopy. The four cell morphologies observed in the presence of *Streptomyces* are depicted: yeast cells, titan cells, hyphal growth, and basidiospore formation. Icons were obtained from BioIcons and modified for this figure. **(B)** Phenotypic profiling of *C. neoformans* following confrontation with *Streptomyces* strains on V8 agar (pH=5) after incubation at room temperature for 4 weeks. Blue circles indicate induction of hyphal development in *C. neoformans* wild-type strains H99α, KN99**a**, and KN99α. PRO, area proximal to *Streptomyces*; DIS, area distal to *Streptomyces*. **(C)** Light microscopy images showing hyphal morphogenesis of *C. neoformans* wild-type strains H99α, KN99**a**, and KN99α following confrontation with *Streptomyces* isolates WAC1490 and WAC1565 on V8 agar (pH=5) after 4 weeks of incubation. Scale bar, 100 µm. See also Figure S1.

As WAC1490 and WAC1565 can grow on diverse media (**Table S1**), we subsequently conducted similar confrontation assays on multiple media (**Figure S1A**). No fungal morphological changes were observed on rich medium YPD or Bennett’s agar, a medium optimized for *Streptomyces* growth, whereas both V8 (pH=5 and pH=7) and MS media supported non-yeast cellular development of *C. neoformans* (**Figure S1B**). Because both V8 and MS media are commonly used to induce sexual reproduction in *C. neoformans*, these results further suggested that the non-yeast cellular development triggered by *Streptomyces* requires conditions conducive to sexual development. Our observations raised the possibility that these morphological changes represent a form of sexual reproduction, such as unisexual reproduction, in *C. neoformans*.

We also found that the effects of *Streptomyces* on *C. neoformans* isolates depend on the proximity between the two species. On MS medium, robust non-yeast development in *C. neoformans* was observed only when cells were positioned within approximately 2.6 cm of *Streptomyces* cells. This distance-dependent effect suggested that diffusible compounds secreted by the bacteria may drive the induction of non-yeast cellular development (**Figure S1C**). Consistent with this hypothesis, in co-culture experiments, morphological changes (primarily the formation of enlarged cells) were first observed on the sides of *C. neoformans* colonies closest to the bacterial cells (after ∼2 weeks). With extended incubation (∼4 weeks), additional development, including robust hyphal growth and sporulation, emerged on the sides of the colonies distal to the bacteria (**Figures 1A**).

Taken together, we identified two natural *Streptomyces* isolates capable of inducing cellular developments resembling unisexual reproduction in wild-type *C. neoformans*, likely through the secretion of diffusible compounds.

### *Streptomyces* induces diverse non-yeast development in *C. neoformans* through secreted tunicamycin

To further characterize the *Streptomyces* isolates analyzed in this study, we generated whole-genome sequencing data for all 31 strains. We then constructed a phylogeny incorporating these isolates along with 15 reference *Streptomyces* species, using concatenated sequences of six housekeeping genes (16S rRNA, *atpD*, *gyrB*, *recA*, *rpoB*, and *trpB*) commonly employed in multi-locus sequence typing (MLST) analyses^41^. The 31 isolates were broadly distributed across the phylogenetic tree.

Notably, the two strains that induced non-yeast development in *Cryptococcus*, WAC1490 and WAC1565, were closely related and clustered within a well-supported clade that also included WAC7446 and two previously reported *Streptomyces* sp. isolates (KY70 and KY75) (**Figure 2A**). These results further suggest that the ability to induce non-yeast development in *Cryptococcus* may be linked to genetic features shared by WAC1490 and WAC1565.

**Figure 2.**
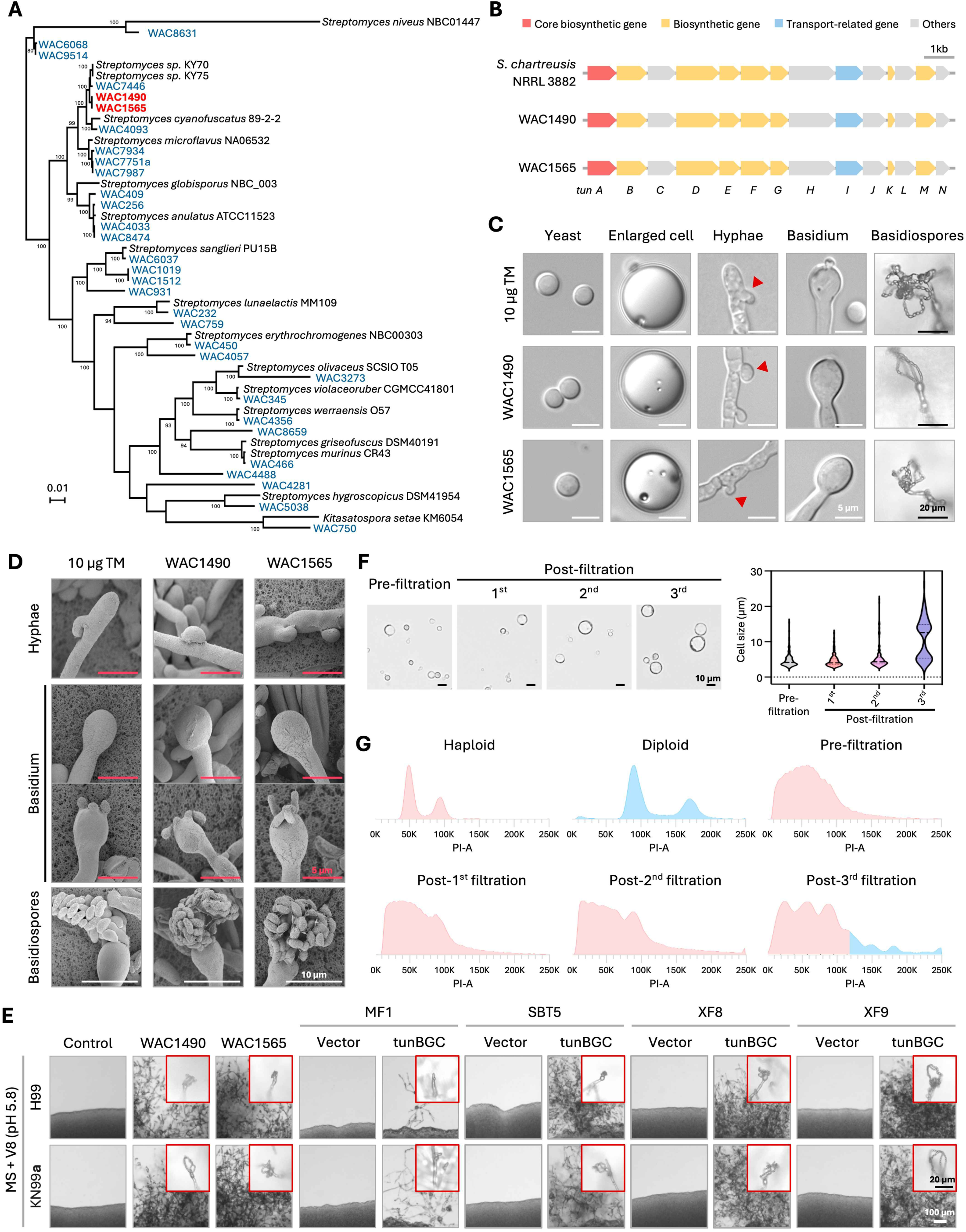
*Streptomyces* isolates drive diverse non-yeast development in *C. neoformans* through secreted tunicamycin. **(A)** Phylogenetic tree showing the relationships of the *Streptomyces* isolates tested in this study (blue and red), as well as several known *Streptomyces* species (highlighted in black). The tree was constructed using concatenated sequences of six conserved genes included in standard MLST analysis, including 16S rRNA, *atpD*, *gyrB*, *recA*, *rpoB*, and *trpB*. Sequences of the *Streptomyces* isolates tested in this study were retrieved from whole-genome sequencing data, and those of the known *Streptomyces* species were retrieved from NCBI. The phylogeny was constructed using the maximum likelihood method implemented in MEGA12^79^, and tested with 1,000 bootstrap replicates. Branches with bootstrap values greater than 85 were highlighted. The two isolates that induced non-yeast development in *Cryptococcus*, WAC1490 and WAC1565, are in red. **(B)** Predicted tunicamycin biosynthetic gene clusters identified in the de novo assembled genomes of *Streptomyces* isolates WAC1490 and WAC1565, using the known tunicamycin BGC in strain *Streptomyces chartreusis* NRRL 3882 as query. **(C)** Light microscopy images of yeast cells, enlarged cells, hyphae, basidia, and basidiospores produced by the *C. neoformans* wild-type strain H99α after confrontation with 10 µg tunicamycin and the two tunicamycin-producing *Streptomyces* isolates. Red arrows along hyphae indicate unfused clamp cells. White scale bar, 5 µm; Black scale bar, 20 µm. **(D)** Scanning electron microscopy images of hyphae, basidium, and basidiospores produced by the *C. neoformans* wild-type strain H99α after confrontation with tunicamycin and the two tunicamycin-producing *Streptomyces* isolates. Red scale bar, 5 µm; White scale bar, 20 µm. **(E)** Confrontation assays between *C. neoformans* wild-type strains (H99α and KN99**a**) and a panel of *Streptomyces* isolates, including the tunicamycin-producing WAC1490 and WAC1565, as well as the four *Streptomyces lividans* isolates (MF1, SBT5, XF8, and XF9) engineered to express the tunicamycin biosynthetic gene cluster (tunBGC) and their respective isogenic empty-vector controls. The confrontation assay was conducted on MS medium supplemented with filtered V8 juice (pH=5.8). Plates were incubated at room temperature in the dark for 4 weeks. Black scale bar, 20 µm; White scale bar, 100 µm. Vector, isogenic strains carrying empty pRT802; tunBGC, *S. lividans* strains carrying pIJ12003a (pRT802::tunBGC). **(F)** Light microscopy images of pre- and post-filtration cell populations of *C. neoformans* wild-type strain H99α, induced for giant cell formation on MS medium supplemented with 5 µg/mL tunicamycin for 5 days. Scale bar, 10 µm. The plot on the right shows the quantification of cell diameters for the pre- and post-filtration populations. **(G)** Flow cytometry analysis of cellular DNA content for the pre- and post-filtration cell populations depicted in panel **(F)**. See also Figures S2 and S3.

Among the *Streptomyces* isolates screened, WAC1490 and WAC1565 are unique in that they both produce tunicamycin, an inhibitor of N-linked glycosylation that can induce endoplasmic reticulum (ER) stress by blocking the GlcNAc phosphotransferase (GPT) enzyme and causing misfolded protein accumulation^42^. We hypothesized that tunicamycin might be the effective metabolite that induces non-yeast development in *C. neoformans*. To test this hypothesis, we first conducted nanopore whole-genome sequencing and assembled the genomes of WAC1490 and WAC1565. We identified a complete tunicamycin biosynthetic gene cluster (tunBGC) in both isolates, with gene organization identical to that of the known tunicamycin-producing *Streptomyces chartreusis* strain NRRL3882 (**Figure 2B**). The tunBGC in WAC1490 and WAC1565 is also identical in sequence and shares ∼97% sequence similarity with the tunBGC in strain NRRL3882. Furthermore, whole-genome sequencing of all screened *Streptomyces* isolates revealed that the tunBGC is only present in WAC1490 and WAC1565, consistent with the observation that only these two isolates induce non-yeast development in *C. neoformans*.

Next, we further tested whether tunicamycin biosynthesis was sufficient to confer the observed fungal morphogenesis rather than by other *Streptomyces* metabolites. To this end, we introduced the tunicamycin biosynthetic gene cluster into the non-tunicamycin-producing species *Streptomyces lividans* MF1 and SBT5^43^, as well as two further optimized derivatives, XF8 and XF9. Each tunBGC-expressing strain was paired to an isogenic empty-vector negative control to distinguish tunBGC-dependent effects from host- or vector-associated effects. LCMS analysis confirmed that tunicamycin production in all heterologous expressing strains, with XF8 and XF9 producing substantially higher levels compared with MF1 and SBT5 (**Figure S3A**). Confrontation assays with *C. neoformans* wild-type strain H99α revealed that, unlike the natural tunicamycin-producing isolates WAC1490 and WAC1565, the heterologous producers did not induce non-yeast development in *C. neoformans* on MS medium (**Figure S3B**); however, they did trigger hyphal development on V8 medium (**Figure S3C**) and induced robust non-yeast development on MS medium supplemented with filtered-V8 juice.

Importantly, the extent of non-yeast development observed in *C. neoformans* was well correlated with the level of tunicamycin produced by the *Streptomyces* isolates (**Figure 2E**): the low-tunicamycin-producing MF1 strain induced only limited hyphal growth and sporulation, whereas the high-tunicamycin-producing isolates XF8 and XF9 elicited robust hyphal growth and sporulation comparable to the morphological changes induced by WAC1490 and WAC1565.

Finally, we asked whether purified tunicamycin alone was sufficient to recapitulate the non-yeast development observed during co-culture with tunicamycin-producing *Streptomyces* isolates. Purified tunicamycin supplementation to MS medium at concentrations ranging from 0.25–1 µg/mL induced non-yeast development, including hyphal growth and sporulation (**Figure S2C**). This was comparable to that observed during co-culture with *Streptomyces* isolates WAC1490 and WAC1565 (**Figures 2C**, **2D and S2B**). However, *C. neoformans* grew only as yeast cells on MS medium with tunicamycin concentrations below 0.25 µg/mL, suggesting that the initiation of non-yeast development requires a minimum level of stress; in contrast, higher concentrations (>2 µg/mL) of tunicamycin repressed sporulation, possibly due to excessive cellular stress.

To further explore potential effects of cellular stress on the observed non-yeast development, we supplemented MS medium with dithiothreitol (DTT), another intracellular ER stress-inducing agent, as well as a broad panel of compounds, including *Streptomyces*-derived antibiotics and antifungals, non–*Streptomyces*-derived antifungals, and mitochondrial respiration inhibitors. None of the test compounds induced non-yeast development in *C. neoformans* wild-type strains (**Figures S2E and S2F**).

Having observed that *C. neoformans* formed enlarged cells in the presence of tunicamycin or tunicamycin-producing *Streptomyces*, we sought to determine whether these cells were titan cells, defined as polyploid cells (>2N) with diameters greater than 10 µm^44^. To test this, we incubated the *C. neoformans* H99α strain on MS medium supplemented with tunicamycin (0.5 µg/mL) and collected cells after 10 days of incubation and prior to the onset of hyphal development. The resulting cell suspension was subjected to three rounds of filtration through a 10 µm pore-size filter, and a subset of cells collected after each round was retained for cell-size estimation and FACS analysis of DNA content. We found that the population became progressively enriched for cells larger than 10 µm in diameter, increasing from 0.26% in the initial population to 54% after the third filtration (**Figure 2F**). Consistent with this, FACS analysis revealed an increased fraction of cells with DNA content exceeding 4C, indicative of polyploidy (**Figure 2G**). Thus, these results support the conclusion that the enlarged cells induced by tunicamycin are indeed polyploid titan cells.

Finally, we assessed the viability of *C. neoformans* yeast cells and basidiospores in the presence of tunicamycin. Our results showed that although yeast cells and spores exhibited similar reductions in survival 24 hours after exposure, the spore population maintained significantly higher viability than yeast cells during prolonged incubation (**Figure S2D**). These findings suggest that the transition to unisexual reproduction and the production of basidiospores may provide a strategy via which *C. neoformans* withstands and escapes exposure to tunicamycin or tunicamycin-producing *Streptomyces*.

Taken together, our findings demonstrate that tunicamycin produced by *Streptomyces* strains is both necessary and sufficient to drive non-yeast cellular development, including processes resembling unisexual reproduction, in *C. neoformans* wild-type strains in a dose-dependent manner. Moreover, this developmental transition may represent an evolutionary adaptation of *C. neoformans* against environmental stresses such as tunicamycin and tunicamycin-producing bacteria.

### *Streptomyces* and tunicamycin induce unisexual reproduction in *C. neoformans*

To further assess whether *C. neoformans* wild-type strains undergo bone fide unisexual reproduction in the presence of tunicamycin or tunicamycin-producing *Streptomyces* isolates, we dissected basidiospores from nine individual basidia produced by the haploid wild-type strain H99α following exposure to tunicamycin (1 basidium), WAC1490 (1 basidium), WAC1565 (4 basidia), and XF8 (3 basidia) (**Table S2**). In addition, we dissected spores from a single basidium produced by the haploid wild-type strain KN99**a** after exposure to WAC1565 (**Table S2**).

Of the 10 basidia analyzed, eight yielded basidiospores with germination rates above 50%, indicating generally high progeny viability consistent with unisexual reproduction (**Table S2**). FACS analysis demonstrated that all of the F1 progeny were haploid (**Figures S4A and S4B**). Importantly, genotyping of the mating-type locus in all of the germinated basidiospores showed that they carried *MAT* alleles identical to those of their parental strain (**Table S2**), supporting the conclusion that these progeny arose through unisexual reproduction involving a single mating type. Additionally, whole-genome sequencing and variant analysis of representative progeny from three H99α basidia and one KN99**a** basidium revealed that basidiospores derived from the same basidium possessed identical genomes that matched those of their parental strains (**Figures 3A and S5A–S5D**).

**Figure 3.**
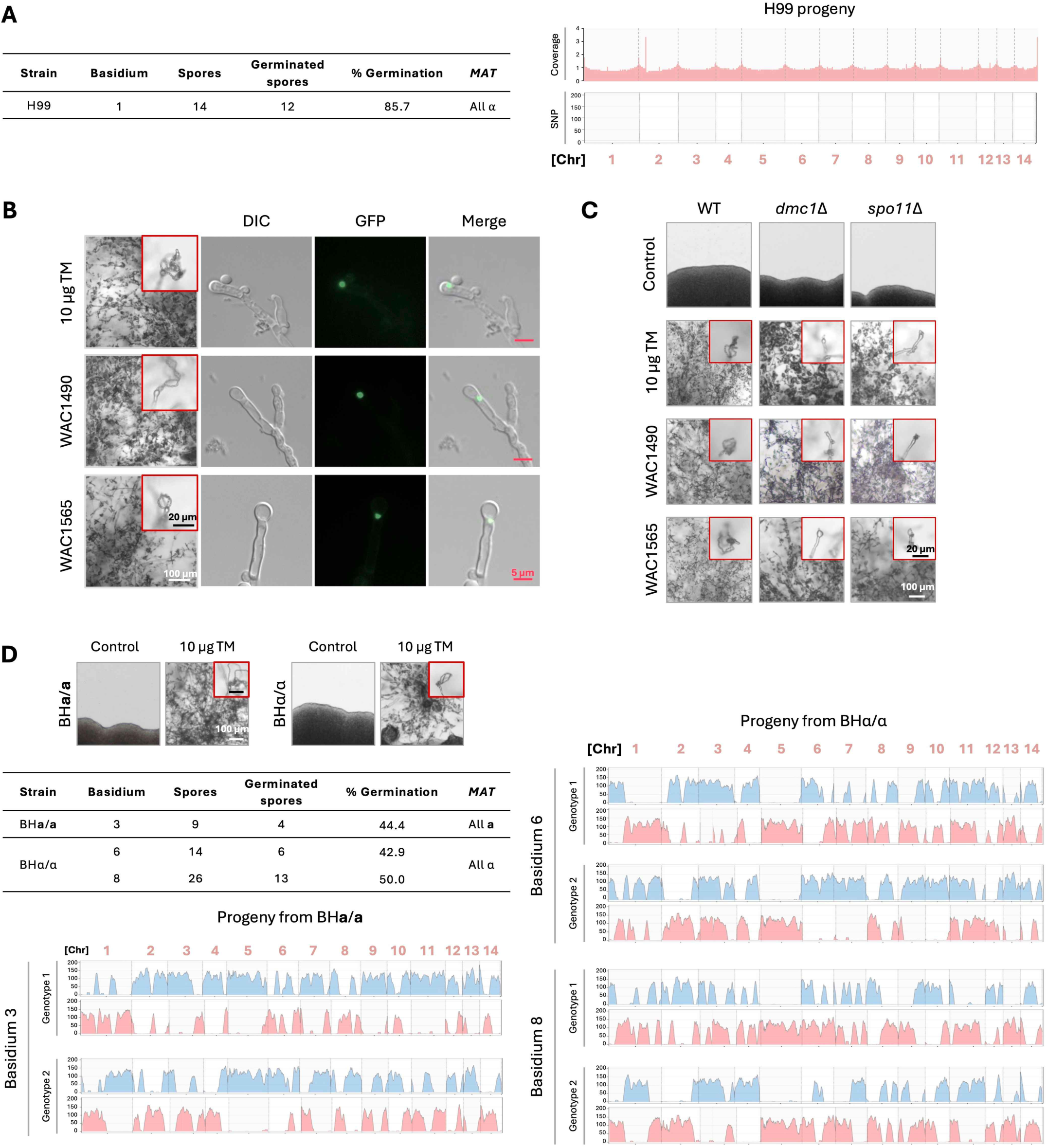
*Streptomyces* and tunicamycin induce unisexual meiotic reproduction in *C. neoformans*. **(A)** Analysis of spores dissected from a single basidium produced by the unisexual reproduction of *C. neoformans* wild-type strain H99α induced by tunicamycin. The table on the left shows the dissection information, including a high spore germination rate and all of the germinated spores being of *MAT*α. The plots on the right illustrate the read coverage (top) and single nucleotide polymorphisms (SNPs, bottom) of one representative progeny using the H99α genome as the reference. Both plots suggest that the progeny has an identical genome as the H99α parental strain. **(B)** Fluorescence microscopy images illustrating the expression of pDmc1-GFP and its localization within the basidia produced by *C. neoformans* wild-type strain H99α cells following confrontation with tunicamycin or with the two tunicamycin-producing *Streptomyces* isolates. Red scale bars, 5 µm; Black scale bars, 20 µm; White scale bar, 100 µm. **(C)** Light microscopy images demonstrating sporulation defects observed in the *C. neoformans* H99α *dmc1*Δ and *spo11*Δ deletion mutants after confrontation with tunicamycin and two tunicamycin-producing *Streptomyces* isolates. The H99α wild-type strain served as a control. Images outlined in red highlight the structures of basidia and basidiospore chains. Black scale bar, 20 µm; White scale bar, 100 µm. **(D)** Analysis of progeny dissected from tunicamycin-induced unisexual reproduction of *C. neoformans* diploid strains BH**a**/**a** or BHα/α, which were fusion products of haploid strains H99α and Bt63**a**, and later underwent loss of heterozygosity at the mating type locus to become *MAT* homozygous. The table lists the dissection information of three individual basidia, one from BH**a**/**a** and two from BHα/α. The spore gemination rates from all three basidia were approximately 50%, suggesting only two of the four meiotic genotypes from each basidium were viable, likely due to genetic divergence between the H99α and Bt63**a** genomes. Plots illustrate the genomic composition of representative progeny from the three basidia, with alleles from Bt63**a** shown in blue and those from H99α depicted in pink. Microscopy images outlined in red highlight the structures of basidia and basidiospore chains. Black scale bar, 20 µm; White scale bar, 100 µm. See also Figures S4–S6.

Taken together, these results provide robust evidence that tunicamycin and tunicamycin-producing *Streptomyces* isolates induce unisexual reproduction in haploid wild-type *C. neoformans* strains.

### Meiosis occurs during unisexual reproduction induced by tunicamycin or tunicamycin-producing *Streptomyces*

Previous studies have shown that unisexual reproduction in *C. neoformans* involves canonical meiosis^37^. To determine whether unisexual reproduction induced by tunicamycin or tunicamycin-producing *Streptomyces* similarly proceeds through meiosis, we employed two complementary approaches.

First, we tested whether two proteins essential for meiosis during *C. neoformans* sexual reproduction, Dmc1 (a meiosis-specific recombinase required for homologous recombination and spore formation) and Spo11 (required for meiotic double-strand break formation and faithful chromosome segregation), are also required for tunicamycin-induced unisexual reproduction^33,45^. To this end, we constructed an H99α strain expressing an ectopic GFP-tagged H3 protein that is placed under the control of the *DMC1* promoter. Exposure of this strain to tunicamycin or the two tunicamycin-producing *Streptomyces* isolates induced robust unisexual reproduction. Fluorescence microscopy revealed basidium-specific expression of Dmc1, indicating that this meiosis-specific protein is expressed within basidia, the sites of meiosis (**Figure 3B**). We also generated H99α strains with deletions of *DMC1* or *SPO11*. Although exposure of these *dmc1*Δ and *spo11*Δ mutants to tunicamycin or tunicamycin-producing *Streptomyces* still induced robust unisexual development, detailed microscopic examination revealed severe sporulation defects. Both mutants predominantly produced basidia with few or no basidiospores (**Figure 3C**). Together, these genetic analyses demonstrate that meiosis-specific genes are activated and required to complete tunicamycin-induced unisexual reproduction, supporting the conclusion that this is a meiotic process.

Second, we directly assessed meiotic recombination employing diploid strains with genome-wide heterozygosity, enabling detection of crossover events. We began with a BH**a**/α strain generated by fusion of the laboratory strain H99α with the natural isolate Bt63**a**, which differs by ∼20,000 SNPs (∼0.1%) across the genome^46^. Because the BH**a**/α strain carries both *MAT***a** and *MAT*α alleles, it is self-fertile. We then screened for BH**a**/α-derived isolates exhibiting markedly reduced self-fertility under standard mating-inducing conditions, indicative of loss of heterozygosity at the *MAT* locus, resulting in *MAT***a**/**a** or *MAT*α/α genotypes. By this approach, we successfully isolated BH**a**/**a** and BHα/α strains.

We next exposed these BH**a**/**a** and BHα/α strains to tunicamycin, which induced robust unisexual reproduction resembling that observed in haploid *C. neoformans* strains (**Figures 3D**). Analysis of spores from four basidia of BH**a**/**a** and eight basidia of BHα/α revealed variable germination rates ranging from 0% to 50% (**Table S2**). The overall lower germination rates, compared to those of the haploid strains, are consistent with the spores being recombinant products derived from genetically divergent H99α and Bt63**a** genomes and are in agreement with previous studies of crosses between these strains^37^. FACS and read-depth analyses of whole-genome sequencing data showed that the vast majority of viable progeny were haploid, with occasional disomy affecting one or two chromosomes (**Figures S4C and S5E**). These findings further support the occurrence of a ploidy-reduction process, such as meiosis, during basidiospore formation.

Finally, variant analysis of the progeny provided clear evidence of allele segregation consistent with meiotic recombination (**Figures 3D and S6**). For example, of the 26 spores dissected from basidium 8 of the BHα/α strain, 13 (50%) germinated. Given that only one round of meiosis occurs within each basidium in *C. neoformans*, producing four recombinant meiotic products^47^, we might expect two distinct nuclear genotypes among the germinated spores at a 50% germination rate. Consistent with this expectation, the 13 viable progeny fell into two genotypic classes, each exhibiting recombinant genomes with alternating segments inherited from the two parental strains along each chromosome, indicative of meiotic crossovers. Similar allele segregation patterns were observed among progeny derived from other basidia of both the BH**a**/**a** and BHα/α strains (**Figures 3D and S6**). These findings are also consistent with the known balanced chromosomal translocation present in the H99 compared with Bt63^48,49^.

Collectively, these analyses demonstrate that tunicamycin-induced unisexual reproduction in *C. neoformans* involves canonical meiotic recombination.

### Tunicamycin-induced unisexual development requires regulators of sexual reproduction and stress response signaling

We next sought to investigate which signaling pathways might underlie tunicamycin-induced unisexual reproduction. We focused on five pathways: 1) pheromone-induced-MAP kinase (MAPK), 2) cAMP/PKA (Protein Kinase A), 3) sexual reproduction-related transcription factors, 4) calcineurin signaling, and 5) HOG (High Osmolarity Glycerol), given their established roles in stress responses and sexual reproduction in *C. neoformans* (**Table S3**) ^50–52^, We obtained deletion mutants for multiple core genes in each pathway in the haploid H99α background and exposed them to tunicamycin or tunicamycin-producing *Streptomyces* to assess effects on non-yeast development (**Figure 4**).

**Figure 4.**
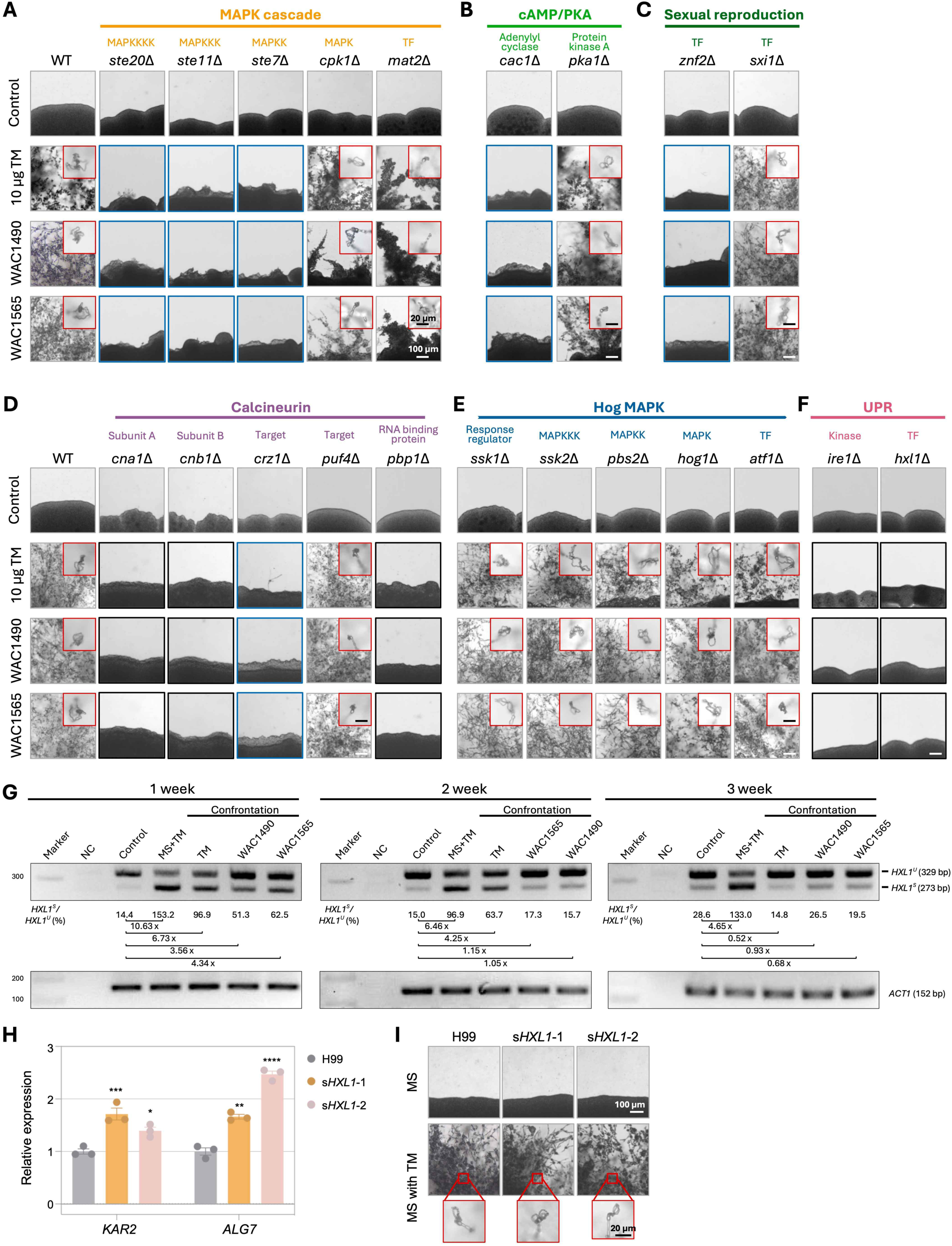
Tunicamycin-induced unisexual development requires regulators of sexual reproduction and stress response signaling. **(A–F)** Light microscopy images of non-yeast development in *C. neoformans* H99α strains in which components of key signaling pathways have been deleted, including those of the MAPK (<u>M</u>itogen-<u>A</u>ctivated <u>P</u>rotein <u>K</u>inase, **A**), cAMP/PKA (<u>P</u>rotein <u>K</u>inase <u>A</u>, **B**), sexual reproduction (**C**), calcineurin (**D**), HOG (<u>H</u>igh <u>O</u>smolarity <u>G</u>lycerol, **E**), and UPR (<u>U</u>nfolded <u>P</u>rotein <u>R</u>esponse, **F**) pathways. Images outlined in blue show production of giant cells without unisexual developments. Images outlined in black show defects in both giant cell production and unisexual developments. Images outlined in red show enlarged light microscopy images highlighting basidiospore formation. Black scale bar, 20 µm; White scale bar, 100 µm. **(G)** Reverse transcription (RT)-PCR analysis of *HXL1* splicing in *C. neoformans* H99α strains following incubation on MS medium containing tunicamycin, or after confrontation with tunicamycin and two tunicamycin-producing *Streptomyces* isolates for 1, 2, and 3 weeks. In each panel, the bottom image shows the *ACT1* control, while the top image displays amplification of the spliced and unspliced *HXL1* alleles, with the numbers underneath indicating the relative quantification of spliced versus unspliced transcripts. H99α strains incubated on MS served as a control in both *ACT1* and *HXL1* RT-PCR. **(H)** Quantitative PCR (qPCR) analysis of the relative expression levels of *HXL1* target genes (*KAR2* and *ALG7*) in *C. neoformans* H99α wild-type strain and two independent engineered strains carrying a spliced allele of *HXL1* (s*HXL1-1* and s*HXL1-2*) integrated into the safe-haven locus. **(I)** Light microscopy images showing cellular developments with (MS with TM) and without (MS) tunicamycin under mating-inducing conditions for *C. neoformans* H99α wild-type strain and strains carrying a spliced allele of *HXL1* integrated into the safe-haven locus. Black scale bar, 20 µm; white scale bar, 100 µm.

Deletion of genes in the HOG pathway did not impair responses to tunicamycin; these strains produced robust hyphae, basidia, and basidiospores, indicating that the HOG pathway is dispensable for tunicamycin-induced unisexual reproduction (**Figure 4E**).

In contrast, deletions in the other four pathways caused defects in unisexual reproduction to varying degrees. Among sexual reproduction-related transcription factors, deletion of *ZNF2* completely abolished tunicamycin-induced unisexual development, whereas deletion of *SXI1*α had no detectable effect (**Figure 4C**).

Similarly, within the cAMP/PKA pathway, deletion of *CAC1* (adenylyl cyclase) abolished unisexual reproduction, while deletion of *PKA1* (catalytic subunit of PKA) did not (**Figure 4B**). These findings indicate that components within the same pathway can contribute differentially to tunicamycin-induced development.

Genes in the calcineurin and MAPK pathways also exhibited diverse roles. In the calcineurin pathway (**Figure 4D**), deletion of *CNA1* (calcineurin A catalytic subunit), *CNB1* (calcineurin B regulatory subunit), *CRZ1* (Calcineurin-responsive zinc finger 1) or *PBP1* (polyadenylated-binding protein 1) completely abolished unisexual development.

The *puf4*Δ mutant displayed responses comparable to the wild type under both conditions, indicating a more limited role in tunicamycin responsiveness. Notably, *puf4*Δ mutants exhibited mild sporulation defects even during unisexual reproduction, suggesting that this gene is required for the efficient completion of this process.

Within the MAPK pathway (**Figure 4A**), deletion of *STE7*, *STE11*, or *STE20* completely abolished tunicamycin-induced unisexual development. The *mat2*Δ mutant showed attenuated responses, with reduced hyphal growth and pronounced sporulation defects. Deletion of *CPK1* did not affect unisexual development in response to tunicamycin alone but reduced hyphal growth during co-culture with tunicamycin-producing *Streptomyces*, suggesting a role in modulating sensitivity to tunicamycin.

Taken together, these results support a model in which tunicamycin responses are regulated by multiple signaling pathways that form a complex, integrated network controlling unisexual development.

### The unfolded protein response (UPR) is essential for tunicamycin-induced unisexual reproduction

Tunicamycin is a well-known inducer of endoplasmic reticulum (ER) stress through inhibition of N-linked glycosylation, leading to the accumulation of misfolded proteins and the perturbation of proteostasis ^42,53^. In *C. neoformans*, ER stress activates the unfolded protein response (UPR), in which the ER-resident sensor Ire1 mediates unconventional splicing of *HXL1* mRNA resulting in excision of a conditional intron and initiation of an adaptive transcriptional program^54^. To determine whether UPR signaling is required for tunicamycin-induced unisexual development, we analyzed *ire1*Δ and *hxl1*Δ deletion strains in the haploid H99α background. Upon exposure to tunicamycin or tunicamycin-producing *Streptomyces*, these UPR-defective strains exhibited a complete loss of tunicamycin-induced unisexual development, indicating that these UPR-related genes are essential for the tunicamycin response (**Figure 4F**).

To gain further insight into the dynamics of the UPR pathway and its potential role in tunicamycin-induced unisexual development, we next performed a quantitative analysis of *HXL1* mRNA splicing in response to tunicamycin. We conducted RT-PCR to assess the two forms of the *HXL1* transcript (i.e., the intron-spliced (UPR-active) and intron-retained (UPR-inactive) isoforms) in cells incubated on MS medium supplemented with tunicamycin, or on MS medium in confrontation with tunicamycin or the tunicamycin-producing *Streptomyces* isolates WAC1490 and WAC1565, for 1, 2, and 3 weeks. These were compared to cells grown on MS medium without tunicamycin (**Figure 4G**).

At all three time points, the highest levels of spliced *HXL1* transcripts were observed in cells grown on tunicamycin-supplemented MS medium, likely reflecting the higher effective concentration of tunicamycin under these conditions. In control samples (MS without tunicamycin), the ratio of intron-spliced to intron-retained *HXL1* transcripts increased over time, suggesting a gradual activation of the UPR pathway, possibly due to accumulating stress during prolonged incubation on nutrient-limited MS medium. In contrast, in cells exposed to tunicamycin, UPR activity peaked after one week of incubation and then gradually declined, as indicated by decreasing fold changes relative to the controls over time (**Figure 4G**). For example, relative to control cells, the ratio of intron-spliced to intron-retained *HXL1* transcripts in cells confronted with tunicamycin decreased progressively from 6.73-fold to 0.52-fold (**Figure 4G**). Combined with the observation that *ire1*Δ and *hxl1*Δ mutants failed to undergo tunicamycin-induced morphological transitions and remained as yeast cells after exposure to tunicamycin (**Figure 4F**), these findings suggest that the UPR pathway plays a critical role at the onset of tunicamycin exposure, facilitating cellular adaptation, developmental remodeling, and the initiation of non-yeast development.

We next constructed two UPR-activated strains in the *C. neoformans* H99α background by inserting an intron-spliced *HXL1* allele (*sHXL1*) into the safe-haven locus. RT-PCR analysis of two target genes known to be induced by the UPR confirmed that the pathway was activated in these modified strains (**Figure 4H**). When grown on MS medium supplemented with tunicamycin, the UPR-activated strains exhibited non-yeast morphology like that observed in the wild-type H99α strain, suggesting that UPR activation did not disrupt the response to tunicamycin. However, when grown on MS medium alone, these strains remained in the yeast form, indicating that activation of the UPR pathway alone was not sufficient to drive the developmental switch from yeast cells to hyphal and unisexual development (**Figure 4I**).

### Transcriptomic profiling reveals temporal dynamics of developmental reprogramming during tunicamycin-induced unisexual reproduction

To gain further insight into how tunicamycin reshapes cellular development in *C. neoformans*, we performed RNA-sequencing analysis with wild-type haploid H99α cells. Samples were collected at two developmental stages following incubation on MS medium supplemented with tunicamycin or two tunicamycin-producing *Streptomyces* isolates: (1) an early stage characterized by robust titan cell formation (2 weeks), and (2) a later stage marked by robust unisexual reproduction (4 weeks).

At the early stage, 244, 182, and 239 differentially expressed genes (DEGs) were identified in cells exposed to tunicamycin, WAC1490, and WAC1565, respectively, relative to controls (**Figure 5A; Table S4**). Among these, 136 genes were shared across all three treatments, including 112 upregulated and 24 downregulated genes.

**Figure 5.**
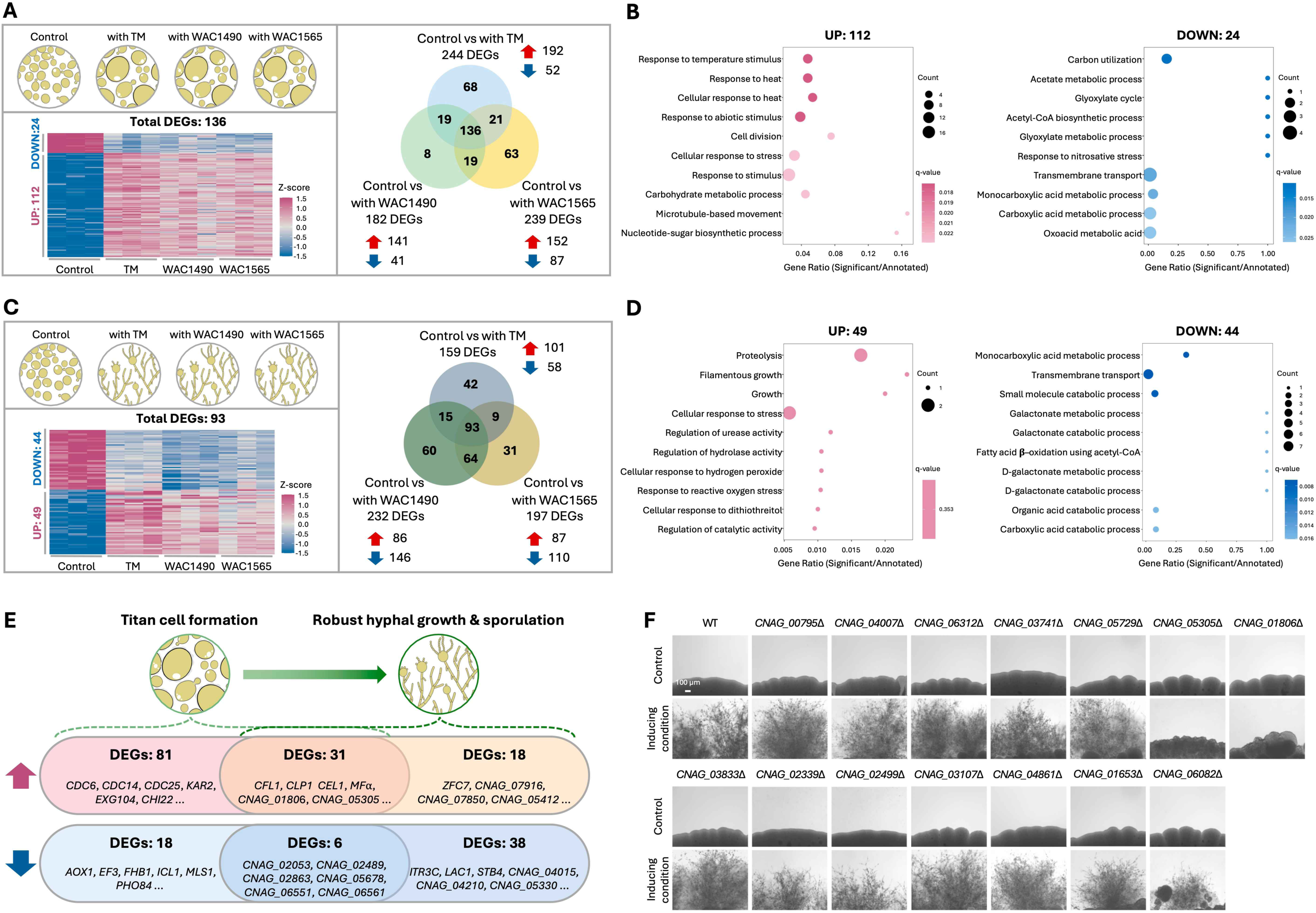
Transcriptomic profiling reveals temporal dynamics of the developmental reprogramming during tunicamycin-induced unisexual reproduction. **(A)** Early-stage differential expression analysis of H99α strains incubated in the presence of tunicamycin or confronted with the two tunicamycin-producing *Streptomyces* isolates (WAC1490 and WAC1565). Cells were collected after two weeks of incubation, at which point the most common non-yeast form development is titan cell formation. H99α cells incubated on the MS medium served as controls. Venn diagram (right) shows shared and unique DEGs, white the heat map (left) depicts the 112 upregulated and 24 downregulated genes shared across all three conditions. **(B)** Gene ontology (GO) enrichment analysis of commonly regulated genes at the early stage (2 weeks). **(C)** Late-stage differential expression analysis of H99α strains incubated in the presence of tunicamycin or confronted with the two tunicamycin-producing *Streptomyces* isolates (WAC1490 and WAC1565). Cells were collected after four weeks of incubation, at which point the non-yeast form developments include titan cell formation, hyphal formation, and sporulation. H99α cells incubated on the MS medium served as controls. Venn diagram (right) shows shared and unique DEGs, while the heat map (left) depicts the 49 upregulated and 44 downregulated genes shared across all three conditions. **(D)** Gene ontology (GO) enrichment analysis of commonly regulated genes at the late stage (4 weeks). **(E)** Stage-specific and overlapping DEGs identified at early and late stages. Numbers indicate upregulated (red) and downregulated (blue) genes in each category. Representative genes are shown. **(F)** Functional analysis of the top 20 early-stage upregulated genes shared among the three conditions. Light microscopy images depict the cellular morphologies of the deletion mutants after four weeks of incubation on MS medium (control) and MS medium supplemented with tunicamycin (unisex-inducing). Deletion of *CNAG_01806* or *CNAG_05305* impaired hyphal development without affecting titan cell formation. Morphological images were created using icons obtained from BioIcons and modified for Figures 5A, 5C, and 5E.

Gene ontology (GO) enrichment analysis revealed that the commonly upregulated genes were associated with responses to heat and other cellular stresses, as expected (**Figure 5B**). More notably, induction of genes involved in cell division and microtubule dynamics suggests engagement of cellular programs associated with ploidy change and developmental differentiation. In contrast, commonly downregulated genes were enriched for carbon utilization pathways, including acetate metabolism and the glyoxylate cycle, as well as transmembrane transport (**Figure 5B**), indicating a coordinated metabolic reprogramming during early adaptation to tunicamycin and morphological changes to titan cell formation.

At the later stage, differential expression analysis identified 159, 232, and 197 DEGs in cells exposed to tunicamycin, WAC1490, and WAC1565, respectively, with 93 DEGs shared across all three treatments, including 49 upregulated and 44 downregulated genes (**Figure 5C; Table S4**). GO analysis of the commonly upregulated genes revealed enrichment for pathways related to proteolysis, filamentous growth, regulation of hydrolase activity, and cellular responses to stress (**Figure 5D**).

Enrichment of filamentation-related genes is consistent with the morphological transition associated with unisexual reproduction. Conversely, commonly downregulated genes were enriched for transmembrane transport and monocarboxylic acid metabolic processes, including galactonate metabolism and fatty acid β-oxidation via acetyl-CoA.

To further delineate transcriptomic changes uniquely associated with early and late responses to tunicamycin, we compared differentially expressed genes identified at these two stages (**Figure 5E**). This analysis revealed 81 and 18 genes that were uniquely up- and downregulated at the early stage, respectively, and 18 and 38 genes that were uniquely up- and downregulated at the later stage. In addition, 31 upregulated and 6 downregulated genes were shared between the two stages (**Figure 5E**).

Among early-stage-specific DEGs, upregulated genes included those involved in the cell cycle, microtubule function, stress responses, and cell wall remodeling, whereas downregulated genes were associated with oxidative responses and translation elongation. In contrast, later-stage-specific upregulated genes were predominantly uncharacterized, while downregulated genes were largely linked to transmembrane transport. For DEGs shared between the two stages, most upregulated genes encoded proteins of unknown function, with a few annotated genes (e.g., *CFL1, CLP1, CEL1*, and mating pheromone genes), whereas downregulated genes were primarily dehydrogenases. Together, these analyses identified both shared and stage-specific transcriptomic changes across treatments and developmental stages induced by tunicamycin.

We observed a greater number of shared up- and downregulated genes among the three treatments (tunicamycin, WAC1490, and WAC1565) at the early stage than at the later stage (**Figures 5A–5D**), suggesting that tunicamycin exerts its most decisive transcriptomic effects early during development. To further elucidate the molecular mechanisms underlying tunicamycin-induced non-yeast development, we therefore focused on early-stage DEGs and identified the top 20 most strongly upregulated genes common to all three treatments (**Table S5**), of which deletion mutants were available for 16 genes^55^.

We next assessed these 16 deletion mutants for their responses to tunicamycin by incubating them on MS medium supplemented with tunicamycin. Deletion of 14 genes exhibited no detectable effect, as these strains still underwent robust non-yeast development, including titan cell formation, hyphal growth, basidium formation, and sporulation (**Figure 5F**). In contrast, deletion of *CNAG_01806* or *CNAG_05305*, encoding two hypothetical proteins, resulted in strains that formed titan cells but failed to undergo hyphal growth, basidium formation, or sporulation (**Figure 5F**). These findings indicate that, while *CNAG_01806* and *CNAG_05305* are dispensable for titan cell formation, they are essential for the yeast-to-hypha transition required for the initiation and completion of tunicamycin-induced unisexual reproduction in *C. neoformans*.

### Natural isolates of *C. neoformans* exhibit varied responses to *Streptomyces* and tunicamycin

Our studies showed that tunicamycin and two tunicamycin-producing *Streptomyces* isolates (WAC1490 and WAC1565) can induce robust unisexual reproduction in *C. neoformans*. We next asked whether this response to tunicamycin is conserved across genetically diverse natural isolates. To address this, we selected a panel of 35 clinical and environmental *C. neoformans* isolates representing the four major lineages: VNI (8), VNII (2), VNBI (11), and VNBII (14). We then assessed their morphological responses to tunicamycin and to the two tunicamycin-producing *Streptomyces* strains on both V8 and MS media (**Figure 6A**).

**Figure 6.**
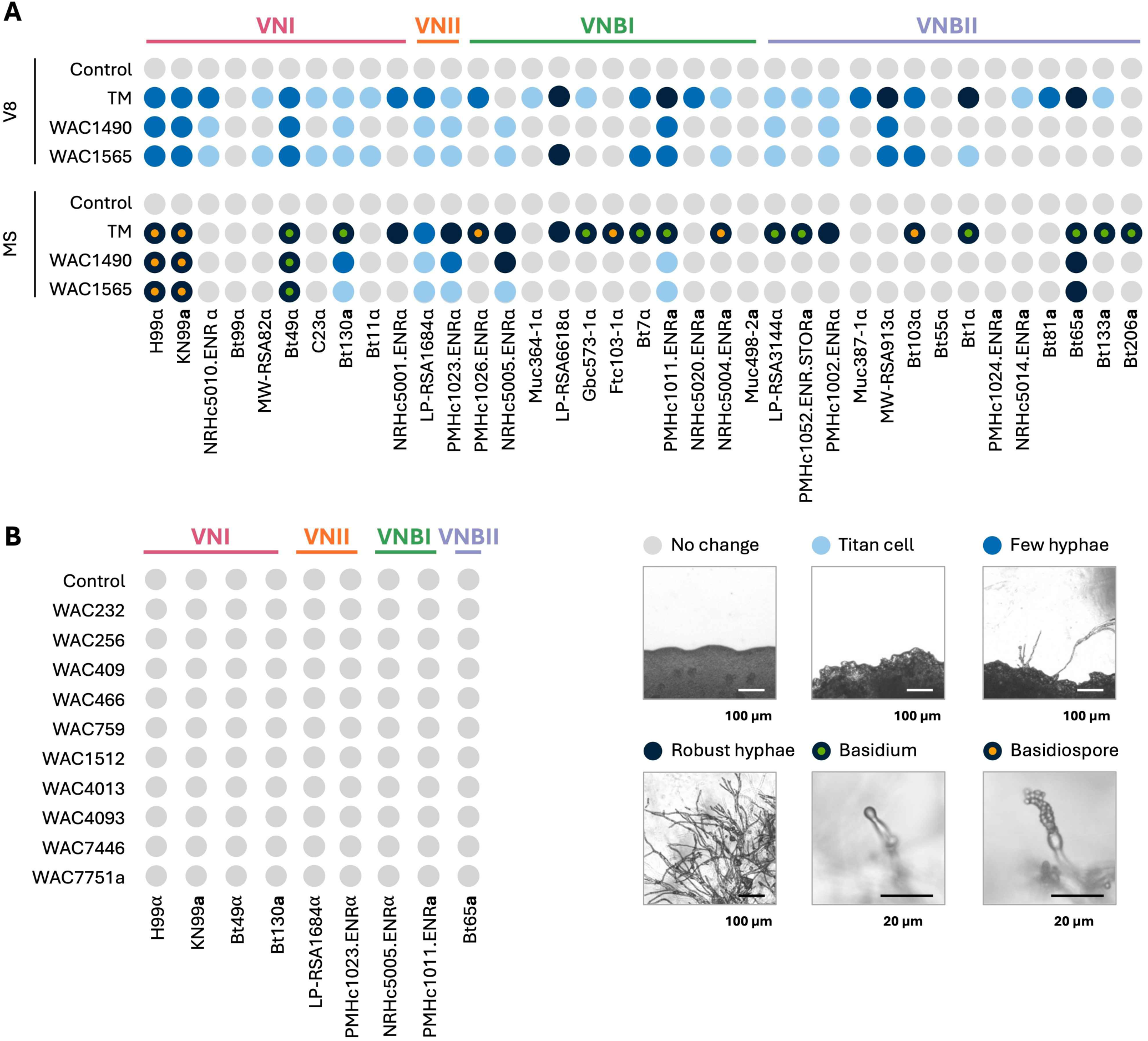
Natural isolates of *C. neoformans* exhibit varied responses to *Streptomyces* and tunicamycin. **(A)** Phenotypic profiling of *C. neoformans* natural isolates following confrontation with tunicamycin and the two tunicamycin-producing *Streptomyces* isolates (WAC1490 and WAC1565) on V8 (pH=5) and MS media plates. The responses of the natural isolates under different conditions are color-coded according to the legends below. **(B)** Phenotypic profiling of *C. neoformans* natural isolates in confrontation assays with ten *Streptomyces* isolates on MS medium plates. For both **A** and **B**, the cellular development observed in the natural isolates are depicted as: no change (grey), titan cell formation (light blue), few hyphae (blue), robust hyphal growth but no basidia (dark blue), hyphal growth with basidia but no spores (dark blue with green dot), and hyphal growth with basidiospore production (dark blue with orange dot).

Thirty-one of the 35 natural isolates exhibited a morphological response to tunicamycin on at least one of the two media tested. The only exceptions were Bt99α (VNI), Muc498-2**a** (VNBI), Bt55α (VNBII), and PMHc1024.ENR**a** (VNBII). These findings suggest that the developmental response to tunicamycin is highly conserved in *C. neoformans*. However, considerable variation in the nature and magnitude of the response was observed among isolates, which was particularly evident when isolates were confronted with the two tunicamycin-producing *Streptomyces* strains on MS medium, where only 7 of 35 *C. neoformans* strains exhibited morphological changes (**Figure 6A**).

In general, stronger responses to tunicamycin, manifested as basidium formation and sporulation, were observed on MS medium. However, a greater number of isolates responded to tunicamycin on V8 medium (29 of 35) compared to MS medium (21 of 35), with some notable exceptions. For example, strains Ftc103-1α (VNBI) and Bt206**a** (VNBII) underwent robust non-yeast development on MS medium but remained in the yeast form on V8 medium. These differences between media suggest additional facets of regulation in the response to tunicamycin, potentially reflecting differences in physiological state that are influenced by both environmental conditions and underlying genetic divergence among *C. neoformans* strains.

Using the seven *C. neoformans* natural isolates that exhibited the most robust tunicamycin-induced unisexual reproduction, we next tested whether other *Streptomyces* isolates could similarly induce non-yeast development (**Figure 6B**). No morphological changes were observed upon exposure to these additional *Streptomyces* strains, further supporting the conclusion that tunicamycin is the primary causative agent driving non-yeast development in *C. neoformans*.

### *Streptomyces* and tunicamycin induce unisexual reproduction in the *C. gattii* species complex

We next sought to determine whether tunicamycin or tunicamycin-producing *Streptomyces* isolates could also induce non-yeast development in the *Cryptococcus gattii* species complex, in which unisexual reproduction remains poorly characterized.

We tested representative isolates from the six species within the *C. gattii* complex and observed robust non-yeast development, including titan cell formation, hyphal growth, and sporulation, in the two congenic *C. deuterogattii* strains, R265**a** and R265α, following exposure to either tunicamycin or tunicamycin-producing *Streptomyces* (**Figures 7A and 7B**). In addition, short hyphae were observed in strains IND107α (*C. tetragattii*) and 7685027α (*C. decagattii*) upon tunicamycin treatment, although neither basidia nor basidiospores were detected in these isolates. To further investigate non-yeast development, we examined the same set of *C. gattii* strains in confrontation assay on MS medium supplemented with filtered V8 juice (MS+V8). Under mating inducing conditions, both R265α and R265**a** strains underwent robust unisexual development, similar to their phenotypes on MS plates. Interestingly, CBS11718**a** (*C. tetragattii*) also underwent complete unisexual reproduction on MS+V8 plates, while WM276α (*C. gattii*) developed short hyphae. These results suggest that the addition of filtered V8 juice further promotes tunicamycin-induced non-yeast development.

**Figure 7.**
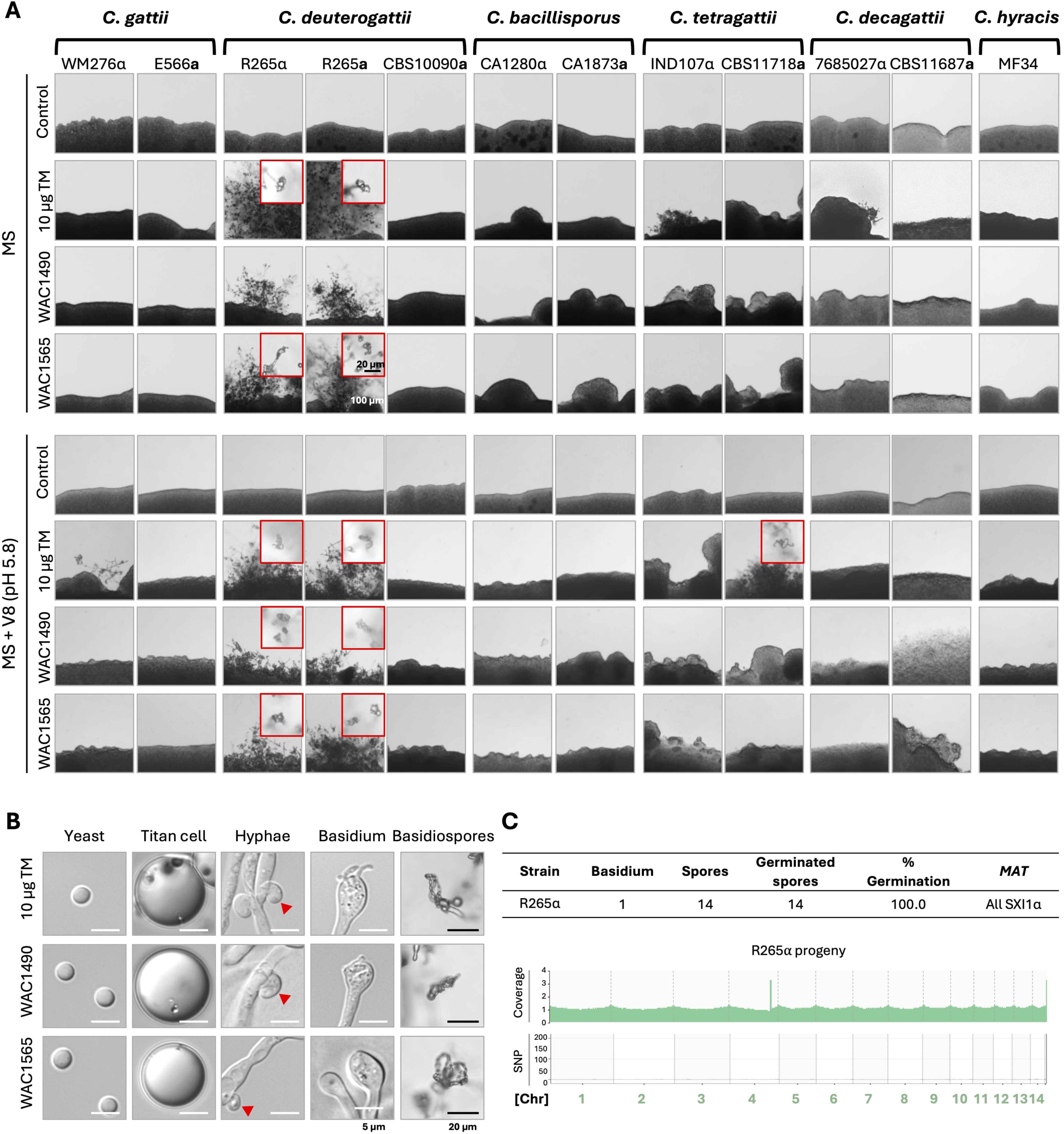
*Streptomyces* and tunicamycin induce unisexual reproduction in the *C. gattii* species complex. **(A)** Light microscopy images of strains from six major lineages of the *C. gattii* species complex following confrontation with tunicamycin and the two tunicamycin-producing *Streptomyces* isolates on MS medium or MS medium supplemented with filtered V8 juice (MS+V8) after incubation at room temperature in the dark for 4 weeks. Images outlined in red highlight the production of basidia and basidiospores by R265α, R265**a,** and CBS11718. Black scale bar, 20 µm; White scale bar, 100 µm. **(B)** Light microscopy images of yeast cells, titan cells, hyphae, basidia, and basidiospores produced by *C. deuterogattii* strain R265α after confrontation with tunicamycin and two *Streptomyces* strains. Red arrows highlight the formation of unfused clamp cells. Red scale bars, 5 µm; Black scale bars, 20 µm. **(C)** Analysis of basidiospores dissected from a single basidium from the unisexual reproduction of the *C. deuterogattii* wild-type strain R265α induced by tunicamycin. The table on top summarizes the spore dissection, highlighting the high germination rate and all of the spores are of the same *MAT*α mating type. The plots below illustrate the read coverage (top) and single nucleotide polymorphisms (SNP, bottom) of one representative progeny using the R265α genome as the reference. Both plots suggest the progeny has an identical genome to the R265α parental strain. See also Figures S7.

We next dissected basidiospores from individual basidia produced during tunicamycin-induced unisexual reproduction in both R265**a** and R265α. These spores exhibited high germination rates, and all viable progeny appeared haploid by FACS analysis, and were of the same mating type as their parental strains (**Figure S7A; Table S6**). Whole-genome sequencing and variant analysis of progeny from the two basidia with the highest germination rates (basidium 1 from R265α and basidium 2 from R265**a**) further demonstrated that their genomes were identical to those of their parents, confirming that they are indeed products of unisexual reproduction (**Figures 7C and S7B–S7C**).

Taken together, these results reveal robust tunicamycin-induced unisexual reproduction in *C. deuterogattii*, highlighting both the conservation of this response across pathogenic *Cryptococcus* species, as well as potential species-specific differences in the trans-kingdom interaction between *Cryptococcus* and *Streptomyces*.

## Discussion

The benefits of sexual reproduction are most commonly attributed to its ability to generate recombinant progeny through meiosis, thereby increasing genotypic and phenotypic variation that can facilitate natural selection and promote adaptation to changing and challenging environments. Previous studies have shown that eukaryotes can adapt their reproductive strategies in response to environmental stresses or the presence of pathogens. For example, freshwater snails switch from inbreeding to outcrossing in the presence of parasites^56–58^. Similarly, *C. elegans* has been shown to mate in response to pathogen exposure^9^. In microorganisms such as fungi, sexual reproduction is often associated with morphological development that confers additional benefits, including filamentous growth that promotes niche expansion and the production of spores that are readily dispersed and more tolerant to environmental stresses^28^.

In this study, we demonstrated that tunicamycin produced by soil-dwelling *Streptomyces* species can induce non-yeast cellular development in the human fungal pathogen *Cryptococcus* species complex, including titan cell formation and unisexual reproduction. Previous studies have shown that titan cells in *Cryptococcus* exhibit elevated resistance to antifungal stresses through mechanisms such as altered cell wall composition and increased ploidy^59^. Additionally, studies in other fungi have demonstrated that hyphal cells are generally more resistant to environmental and antifungal stresses than yeast cells^60^. Furthermore, as shown in our study, basidiospores produced through unisexual reproduction display greater long-term resistance to tunicamycin than yeast cells. Together, these findings suggest that the developmental responses we observed may represent evolutionary outcomes of persistent inter-kingdom interactions between *Cryptococcus* and antimicrobial-producing microbial co-inhabitants in natural environments. Such stresses impose selective pressures that drive the evolution of traits enabling *Cryptococcus* either to withstand these threats (e.g., production of titan cells, hyphae, and dormant spores) or to escape them through dispersal mechanisms such as hyphal expansion and spore production.

It has been hypothesized that titan cell formation in *Cryptococcus* evolved in response to interactions with amoebae in soil environments, and both titan cells and basidiospores are known to play critical roles in *Cryptococcus* pathogenesis ^61^. Our study thus suggests that these medically important traits may also represent byproducts of selective pressures imposed by antimicrobial compounds produced by cohabiting bacterial species, such as *Streptomyces*, in shared environmental niches. Our observation that tunicamycin induced unisexual reproduction in some, but not all, lineages within the *C. gattii* species complex suggests that the response to tunicamycin-like antimicrobials is evolutionarily conserved but exhibits naturally occurring variation. This variation may reflect differences either in the ability of *Cryptococcus* species to respond to these compounds or in the specific environmental cues to which distinct lineages are programmed to respond.

With the exception of the Hog MAPK pathway, all of the pathways we analyzed exhibited at least partial involvement in *C. neoformans* responses to tunicamycin and tunicamycin-producing bacteria, highlighting the complex regulatory networks underlying titan cell formation and sexual development in *Cryptococcus*^62^. We observed variation in the responses of different deletion mutants to tunicamycin, with some mutants defective in both titan cell formation and unisexual development, whereas others retained the ability to form titan cells but failed to undergo unisexual reproduction. These observations are consistent with the finding that, in response to tunicamycin and tunicamycin-producing bacteria, titan cell formation consistently preceded unisexual development in *Cryptococcus*, further supporting the proposed interconnection between these two cellular programs^32^.

Recent studies have shown that unisexual reproduction in *C. neoformans* and *C. deneoformans* can be induced by deletion of the *RIC8* gene, which encodes a guanine nucleotide exchange factor (GEF) that modulates G-protein activity^32,37^. However, we did not observe pronounced differences in *RIC8* expression in our transcriptomic analyses. This finding may suggest that tunicamycin-induced unisexual reproduction is regulated through pathways distinct from those previously described. Alternatively, the requirement for Ric8 down-regulation for cellular development reprogramming may be transient or occur at an early stage that is prior to our first sampling time point (2 weeks). It is also possible that tunicamycin may reduce Ric8 protein abundance or activity through post-transcriptional mechanisms that would not be captured at the transcript level. One of the genes highly upregulated in our transcriptomic analysis was *CFL1*, a key adhesion protein in *C. neoformans* that coordinates the yeast-to-hyphae transition through paracrine signaling^63^. The upregulation of *CFL1* was observed in both early and late stages of our transcriptomic datasets, suggesting that tunicamycin may promote sustained activation of developmental pathways associated with *CFL1*.

Tunicamycin is a canonical inducer of ER stress and, in our study, acted as a potent inducer of titan cell formation and unisexual reproduction in *Cryptococcus*. However, another ER stress–inducing agent, DTT, failed to trigger comparable developmental transitions under our experimental conditions. This suggests that generalized ER stress alone may be insufficient to induce these developmental programs and that specific cellular consequences of tunicamycin exposure, such as inhibition of N-linked glycosylation or selective activation of downstream signaling pathways, may be required. However, we cannot rule out the possibility that the experimental conditions tested, including media composition and growth environment, were not optimal for eliciting similar responses to DTT.

Sexual reproduction in the choanoflagellate *Salpingoeca rosettaa* can be stimulated by a factor released by a cohabiting bacterium^8^. Similarly, we demonstrate here that a defined compound produced by a natural microbial co-inhabitant can induce pathogenicity-related traits, including titan cell formation and unisexual reproduction, in a medically important opportunistic human fungal pathogen. Sex is considered universal and plays a critical role in shaping the population structure and evolutionary trajectory of a species. While comparative and population genetic/genomic studies have suggested that sexual reproduction is widespread among unicellular eukaryotes such as fungi, direct observation of sexual cycles under laboratory conditions remains challenging for many species, including some human fungal pathogens that appear to have a clonal population with highly skewed mating-type compositions, such as different lineages in the *Cryptococcus* species complexes and *Candida auris*. In addition to the myriad environmental factors known to influence fungal sexual reproduction (e.g., temperature, moisture, nutrient availability, and light, etc.), biotic interactions with co-inhabitants in their native ecological niches may represent critical cues governing these otherwise cryptic sexual cycles in nature.

## STAR Methods

### Fungal strains, media and growth conditions

*Cryptococcus* and *Streptomyces* strains used in this study are listed in **Table S7** and **Table S8**. *Streptomyces* strains were grown on Bennett’s medium (soluble starch, 10 g/L; casamino acids, 2 g/L; yeast extract, 1.8 g/L; Czapek mineral mix, 2 mL; agar, 20 g/L). The Czapek mineral mix contained 10 g KCl, 10 g MgSO₄·7H₂O, 12 g NaNO₃, 0.2 g FeSO₄·7H₂O, and 200 µL concentrated HCl, adjusted to 100 mL with double-distilled water. *Cryptococcus* strains were generally grown in liquid or on solid yeast extract peptone dextrose (YPD; BD, Franklin Lakes, NJ, USA) medium at 30°C. To induce sexual development, *Cryptococcus* strains were incubated on solid V8 medium (V8 juice, 50 mL; KH₂PO₄, 0.5 g/L; agar, 40 g/L) or Murashige and Skoog medium (MS powder, 4.328 g/L; agar, 20 g/L; pH adjusted to 5.8) and incubated at room temperature (22–25°C) in the dark^34^.

To generate strains carrying spliced *HXL1* integrated at the safe-haven locus, an approximately 2 kb fragment containing the *HXL1* promoter and the 5′ portion of the ORF, along with a 1 kb fragment containing the terminator region, was amplified using primers listed in **Table S9**. These fragments were designed to exclude the unconventional *HXL1* intron and were cloned into the pSDMA57 vector^64^. The resulting plasmid was digested with PacI and co-transformed into the *C. neoformans* H99 background along with the corresponding gRNAs and Cas9 expression cassette.

### Confrontation assay

Single colony of *Cryptococcus* strains were inoculated into liquid YPD medium and cultured at 30°C with shaking at 80 rpm for 16–18h. Approximately 10^7^ spores of *Streptomyces* were inoculated in liquid YPD medium and cultured under same conditions. For confrontation assays, 5 μL of the *Streptomyces* culture was spotted at the center of solid YPD, Bennett’s, V8 (pH=5 or pH=7), or MS medium. *Cryptococcus* cells were harvested, washed twice with distilled water, and adjusted to an OD_600_ of 0.1. A 5 μL aliquot of the *Cryptococcus* suspension was then spotted 1.3 cm away from the *Streptomyces* spot. Plates were incubated at room temperature in the dark for approximately 4 weeks. Colony morphology and hyphal development were examined using a Zeiss light microscope equipped with an Axiocam 105 color camera (ZEISS, Oberkochen, Germany)^34,65^. All confrontation assays were performed at least twice independently. Two independent deletion mutants were tested for each gene involved in signaling pathways.

### Viability assessment of *C. neoformans* yeast cells and basidiospores

Basidiospores were dissected from H99α × KN99**a** crosses grown on MS medium for 1 week. For controls, yeast cells of H99α and KN99**a** incubated on the same MS medium were harvested at the same time as the basidiospores. Both basidiospores and yeast cells were resuspended in PBS containing tunicamycin (5 μg/mL) at a density of ∼5 cell/μl. After 0, 1, 2, and 3 days of incubation, 200 ul of cell suspension were spread onto YPD agar plates, and cell viability were estimated based on the colony-forming units after incubation at 30 °C for 3 days.

### Scanning electron microscopy (SEM)

Hyphae and unisexual structures formed during confrontation assays were first examined by light microscopy. Agar blocks containing the structures were excised and fixed in 4% (v/v) formaldehyde (Sigma) and 4% (v/v) glutaraldehyde (Sigma) at 4°C overnight without agitation. The fixative was removed, and samples were serially dehydrated in 30%, 50%, 70%, and 95% (v/v) ethanol, with 1h incubation at 4°C at each step. Samples were then washed three times in 100% ethanol at room temperature, dried using a Ladd CPD3 critical point dryer (Ladd Research Industries, Essex Junction, VT, USA), and sputter-coated with gold using a Denton Desk V coater (Denton Vacuum, Moorestown, NJ, USA). Hyphae with unfused clamp cells, basidia, and basidiospores were imaged with a scanning electron microscope equipped with an EDS detector (Apreo, ThermoFisher)^66^.

### Fluorescence-activated cell sorting (FACS) analysis

To determine the ploidy of progeny derived from wild-type haploid *Cryptococcus*, cells were grown on solid YPD medium for two days and then harvested. Cells were washed twice with PBS and fixed in 70% ethanol at 4°C overnight with agitation. The following day, the ethanol was removed, and the cells were washed once with 500 μL of 1x nuclear staining (NS) buffer (10 mM Tris-HCl, pH 7.5; 25 mM sucrose; 0.95 mM MgCl₂; 0.1 mM CaCl₂; 0.1 mM ZnCl₂; 1 mM EDTA, pH 8.0; 0.6 mM PMSF; 7 mM 2-mercaptoethanol). Samples were resuspended in NS buffer containing propidium iodide (PI; Sigma) and incubated at room temperature in the dark for at least 2h. Subsequently, 50 μL of the stained cell suspension was mixed with 500 μL of Tris-PI solution, and DNA content was analyzed by FACS analysis at the Duke Cancer Institute Flow Cytometry Shared Resource Laboratory. Data were analyzed using FlowJo (v10.10.0) software^67^.

### Analysis of titan cells induced by tunicamycin

To determine the ploidy of giant cells induced by tunicamycin, *Cryptococcus* cells were spread onto MS medium supplemented with 5 µg/mL of tunicamycin and incubated at room temperature for 10 days. A mixture of yeast and giant cells was harvested, resuspended in PBS, and filtered three times with a 10 µm pluriStrainer (pluriStrainer, USA, El Cajon, CA, USA) to enrich for large cells. The collected cells were examined by light microscopy, and cell size was quantified using ImageJ and plotted with GraphPad Prism v10.1.1. Cells were then pelleted by centrifugation, washed with 70% ethanol, and subjected to FACS analysis to assess DNA content and polyploidy.

### Heterologous expression of the tunicamycin gene cluster

The tunicamycin biosynthetic gene cluster (*tun* BGC) was cloned into the SacI site of pRT802 to generate pIJ12003a^68^. The empty pRT802 vector served as a negative control. Plasmids pIJ12003a and pRT802 were first propagated in *E. coli* DH5α and subsequently introduced into the engineered background-cleaned *Streptomyces lividans* host strains SBT5, MF1, XF8, and XF9 by triparental intergeneric conjugation using *E. coli* ET12567/pR9406 as the helper strain^43,68^. Kanamycin-resistant exconjugants were selected on mannitol soya flour (2% mannitol, 2% soya flour, and 2% agar) plates containing trimethoprim and kanamycin.

To assess heterologous tunicamycin production, recombinant *S. lividans* strains harboring either pIJ12003a or the empty pRT802 vector were cultivated and their fermentation extracts were analyzed by LC-HRMS/MS. Strains were first grown on mannitol soya flour agar medium at 30°C for 7 days to promote sporulation. A single colony or sporulated culture was then transferred to Bennett’s agar plates and incubated at 30°C for an additional 7 days for metabolite production.

### Extraction and LC-HRMS/MS analysis of tunicamycin production

Fermentation cultures were extracted with methanol. The methanolic extracts were dried under reduced pressure using a Genevac evaporator (Canadawide Scientific), and the residues were resuspended in DMSO prior to LC-HRMS/MS analysis.

HRMS analyses were performed on an Agilent 1290 Infinity II LC system coupled to an Agilent 6550 qTOF mass detector. The instrument was operated in positive ionization mode with the following source parameters: capillary voltage, 3500 V; nozzle voltage, 1000 V; fragmentor voltage, 380 V; dry gas flow rate, 14 L/min; dry gas temperature, 200°C; nebulizer pressure, 35 psig; sheath gas temperature, 350°C; and sheath gas flow rate, 11 L/min. Data were acquired over an m/z range of 100–3000 at a rate of 1 spectrum/s.

Chromatographic separation was performed on an Eclipse SDB-C8 column (2.1 mm × 100 mm, 3.5 µm; Agilent) at a flow rate of 0.4 mL/min. Mobile phase A was H₂O with 0.1% formic acid, and mobile phase B was acetonitrile with 0.1% formic acid. The gradient started at 25% B for 0.5 min, followed by a linear increase to 100% B over 6.5min.

Tunicamycins were identified by comparison with a commercial tunicamycin standard mixture (T7765-5MG) and by HRMS/MS-based GNPS molecular networking. For GNPS analysis, raw HRMS/MS data were converted to mzXML format and analyzed using the GNPS online workflow. Networks were generated using precursor and fragment ion tolerances of 1.0 Da and 0.5 Da, respectively, with edges requiring a cosine score >0.7 and at least six matched peaks. Spectral library matches were accepted using the same criteria^7,69^.

### Quantification of tunicamycin production in *S. lividans*

Tunicamycin production in recombinant *S. lividans* strains was quantified by LC-MS using calibration curves generated from a commercial tunicamycin standard mixture. Extracted ion chromatograms were generated for tunicamycin C ([M+H]⁺ m/z 817.41), tunicamycin V ([M+H]⁺ m/z 831.42), tunicamycin B ([M+H]⁺ m/z 845.44), and tunicamycin D ([M+H]⁺ m/z 859.46). Calibration curves were established by plotting peak area against standard concentration across a six-point dilution series ranging from 12.5 to 800 µg/mL.

### Sexual reproduction and spore dissection

To analyze progeny produced by *Cryptococcus* solo cultures in confrontation with *Streptomyces* or in the presence of tunicamycin, basidium-specific basidiospores were microdissected as previously described^34^.

### Nanopore sequencing and genome assembly of *Streptomyces* isolates

Two *Streptomyces* isolates were cultured overnight in liquid YPD medium at 30°C and harvested by centrifugation. Cells were pre-lysed in STET buffer (10 mM Tris-HCl, pH 8.0, 0.1 M NaCl, 1 mM EDTA, and 5% Triton X-100) supplemented with lysozyme (1 mg/mL; Sigma) at 37°C for 30 min. High molecular weight (HMW) DNA was extracted using the Monarch HMW DNA Extraction Kit for Tissue (New England Biolabs, Ipswich, MA, USA). DNA quality and quantity were assessed using a Nanodrop (ThermoFisher) and a Qubit Fluorometer with a Qubit dsDNA High Sensitivity kit (ThermoFisher). DNA integrity was confirmed by a pulsed-field gel electrophoresis, revealing intact HMW DNA. Long-read sequencing was performed by Plasmidsaurus (South San Francisco, CA, USA) using Oxford Nanopore Technologies (ONT; Oxford, United Kingdom). Libraries were prepared using v14 chemistry and sequenced on R10.4.1 flow cells. Raw reads were error-corrected with Canu v2.2^70^, followed by assembly with Flye v2.9.5-b1801 using parameters specifying an expected genome size of 10 Mb and a minimum overlap length of 5 kb^71^. Assemblies were polished using Pilon (v1.24) with default parameters and Illumina short-read data. Gene annotation was performed using Bakta (v1.9.4). This pipeline generated high-quality genome assemblies: WAC1490 genome assembled into two contigs (N50 = 111.8 kb; 99.8% BUSCO completeness; 7,313 predicted genes), and WAC1565 genome assembled into two contigs (N50 = 96.9 kb; 98.8% BUSCO completeness; 7,314 predicted genes)^72,73^.

### Whole-genome Illumina sequencing and small nucleotide polymorphism (SNP) analysis

*Cryptococcus* strains were cultured on solid YPD medium at 30°C, harvested, and lyophilized. Lyophilized cells were mechanically disrupted, mixed with lysis buffer (Lucigen, Middleton, WI, USA), and incubated at 65°C for 45 min. After cooling on ice, samples were centrifuged, and the supernatants were transferred to new tubes and mixed with MPC Protein Precipitation Reagent (Lucigen). Following centrifugation, the clarified supernatants were transferred to new tubes, mixed with an equal volume of isopropanol (ThermoFisher), and centrifuged to pellet genomic DNA. The supernatant was discarded, and the DNA pellet was washed with 70% ethanol, air-dried, and resuspended in 1x EB buffer (Qiagen) for 30min. DNA quality and quantity were assessed using a Nanodrop (ThermoFisher) and a Qubit Fluorometer with the Qubit dsDNA High Sensitivity kit (ThermoFisher).

Illumina sequencing was performed at the Duke Sequencing and Genomic Technology core facility. Genomic DNA libraries were prepared using the KAPA HyperPrep Kit (Roche, Basel, Switzerland) and paired-end sequencing (2 × 151 bp) was conducted on the NovaSeq X platform (Illumina, San Diego, CA, USA). Raw FASTQ files were quality trimmed using Trimmomatic (v0.39)^74^. Trimmed reads were aligned to the reference genomes of *C. neoformans* H99 (GenBank: GCA_000149245.3) or *C. deuterogattii* R265 (GenBank: GCA_002954075.1) using Snippy (v4.6.0, RRID:SCR_023572). VCF calling was performed relative to the parental reference datasets, including *C. neoformans* H99α (SRR647805) and *C. deuterogattii* R265 (SRR643936). SNP density was calculated as the number of SNPs per 10 kb genomic window, and results were visualized using RStudio (v4.3.2).

### RNA extraction, reverse transcription PCR (RT-PCR), and quantitative PCR (qPCR) analysis

To isolate total RNA from the *Cryptococcus* cells following confrontation, cells were harvested, suspended in PBS, and centrifuged. Collected cells were lyophilized overnight and homogenized using a FastPrep bead-beater (MP Biomedicals, Santa Ana, CA, USA) with 0.2 mL of zirconia-silica beads (BioSpec Products, Bartlesville, OK, USA). Total RNA was purified with the RNeasy PowerPlant Kit (Qiagen, Hilden, Germany). RNA quality and quantity were analyzed using a Nanodrop (ThermoFisher, Waltham, MA, USA) and a Qubit Fluorometer with a Qubit RNA High Sensitivity kit (ThermoFisher)^75^. Following DNase treatment, cDNA was synthesized using the Maxima H Minus cDNA Synthesis Master Mix (ThermoFisher). To evaluate *HXL1* splicing, RT-PCR was performed using the primers listed in **Table S9**, with *ACT1* serving as an internal control. For qPCR analysis, reactions were set up in technical triplicate using PowerUp SYBR Green Master Mix (ThermoFisher) on a QuantStudio 3 qPCR system (ThermoFisher). Primers for qPCR are listed in **Table S9**. Gene expression was normalized to *ACT1*, and relative expression was calculated using the 2^−ΔΔCt^ method.

### Transcriptomic analyses

*C. neoformans* wild-type H99 cells were grown on MS medium either alone (control) or in the presence of tunicamycin or two *Streptomyces* isolates (WAC1490 and WAC1565). Samples were collected at two developmental stages: an early stage (2 weeks after incubation, following titan cell formation but prior to hyphal differentiation) and a late stage (4 weeks after incubation, following hyphal formation and sporulation). Scraped cells were lyophilized overnight, and total RNA was extracted as described above. RNA samples were submitted for sequencing at the Duke University Sequencing and Genomic Technologies core facility, and libraries were prepared with the Watchmaker mRNA library kit and sequenced on an Illumina NovaSeq X Plus platform to produce 150-bp paired-end reads.

Raw reads were trimmed using Trimmomatic (v0.39)^74^, and trimmed reads were mapped to the reference genome of *C. neoformans* H99 (GenBank: GCA_000149245.3) using the STAR RNA-seq aligner (v2.7.11b)^76^. Gene-level counts were generated from aligned BAM files using featureCounts (Subread v2.1.1)^77^.

Differentially expression analyses were performed using DESeq2 (v1.40.2)^78^, and genes with |log₂ fold change| ≥ 1 and false discovery rate (FDR) < 0.05 were considered differentially expressed. Data visualization was performed using the R package ggplot2 and GO enrichment analysis was conducted using topGO. Results were visualized in RStudio (v4.3.2). All RNA-seq experiments were performed with three biological replicates.

## Data availability

Whole-genome sequencing raw data and RNA sequencing raw data are available in the NCBI Sequence Read Archive (SRA) under PRJNA1455727.

## Acknowledgments

We are grateful to Dr. Paul Magwene and Deborah Murray for sharing preliminary observations about tunicamycin inducing complex colony morphology on YPD medium including formation of hyphae. We appreciate Dr. Vikas Yadav, Connor Larmore, and Anna Lehmann for critically reading the manuscript. This study was supported by the NIH/National Institute of Allergy and Infectious Diseases (R01-AI039115-28, R01-AI050113-20, R01-AI170543-05, and R01-AI172451-04) awarded to J.H. and the Canadian Institutes for Health Research (PJT190298) to G.W.. J.H. is co-director and fellow and G.W. is fellow of the CIFAR program Fungal Kingdom: Threats & Opportunities.

## Author contributions

Conceptualization: Y.S., S.S. and J.H.; Methodology: Y.S. and S.S.; Software: Y.S.; Validation: Y.S. and S.S.; Investigation: Y.S., S.S. and X.C.; Visualization: Y.S.; Resources: J.H.; Writing – Original Draft: Y.S. and S.S.; Writing – Review & Editing: Y.S., S.S., X.C., G.W. and J.H.; Supervision: S.S. and J.H.; Funding Acquisition: J.H.

