## Supplementary Figures and Tables for "*Streptomyces* induces unisexual reproduction of a major human fungal pathogen"

**Supplemental information**

Figures S1–S7, and Tables S1–S9

Figure S1. Two *Streptomyces* isolates induce diverse non-yeast development in *C. neoformans* strain H99α, related to Figure 1.

Figure S2. *Streptomyces* isolates drive developmental reprogramming in *C. neoformans*, related to Figure 2.

Figure S3. Tunicamycin is the causative agent for the induction of unisexual reproduction in *C. neoformans*, related to Figure 2.

Figure S4. Ploidy analysis of basidiospores produced by unisexual reproduction of *C. neoformans* wild-type haploid (KN99**a** and H99α) and diploid (BH**a**/**a** and BHα/α) strains, related to Figure 3.

Figure S5. WGS and variant analysis of basidiospores produced from *Streptomyces*- and tunicamycin-induced unisexual reproduction in *C. neoformans* wild-type haploid strains (KN99**a** and H99α) and diploid strains (BH**a**/**a** and BHα/α), related to Figure 3.

Figure S6. *Streptomyces*- and tunicamycin-induced unisexual reproduction in *C. neoformans* involved meiotic recombination, related to Figure 3.

Figure S7. Analyses of basidiospores confirmed that *Streptomyces* and tunicamycin induced unisexual reproduction in *C. deuterogattii*, related to Figure 7.

Table S1. Assessment of bacterial growth on various culture media, related to Figure 1.

Table S2. Genotypic analysis of unisexual spores produced from *Cryptococcus*. *neoformans* wild-type haploid strains KN99**a** and H99α and diploid strains BH**a**/**a** and BHα/α, related to Figure 3.

Table S3. Summary of genes involved in sexual reproduction and stress response signaling pathways in *C. neoformans*, related to Figure 4.

Table S4. Commonly regulated differentially expressed genes at early and/or late stages following treatment with tunicamycin or exposure to two tunicamycin-producing *Streptomyces* isolates (WAC1490 and WAC1565), related to Figure 5.

Table S5. Top 20 differentially up-expressed genes after titan cell formation, related to

Figure 5.

Table S6. Genotypic analysis of unisexual spores produced from *Cryptococcus deuterogattii* wild-type R265**a** or R265α, related to Figure 7.

Table S7. *Streptomyces* strains used for confrontation assay.

Table S8. *Cryptococcus* strains analyzed in this study.

Table S9. Oligonucleotides utilized for this study.

**Figure S1. Two *Streptomyces* isolates induce diverse non-yeast development in *C. neoformans* strain H99α, related to Figure 1.**

**(A)** The induction of diverse non-yeast development in the *C. neoformans* wild-type strain H99α by the two *Streptomyces* isolates (WAC1490 and WAC1565) were tested on various media, including YPD, Bennett's, and V8 (pH=7), with V8 (pH=5) and MS agar media serving as controls. No non-yeast form development was observed on YPD and Bennett's medium plates. **(B)** Light microscopy images of hyphal morphogenesis in *C. neoformans* wild-type strain H99α following confrontation with the two *Streptomyces* isolates on V8 (pH=5 and pH=7) and MS medium plates. Scale bar, 100 µm. **(C)** Distance-dependent induction of non-yeast development in *C. neoformans* wild-type strain H99α by the two *Streptomyces* isolates (WAC1490 and WAC1565). The assay was conducted on MS medium plates, with bacterial and *C. neoformans* cells spotted at varying distances (1, 1.5 and 2 units, with 1 unit = 1.3 cm). After 4 weeks of incubation, on the side of *C. neoformans* spots that are distal to the bacterial cells, robust non-yeast developments were observed on the 1-unit plate, no non-yeast development was observed on the 2-unit plate, and intermediate levels of non-yeast developments were observed on the 1.5-unit plate.

**Figure S2. *Streptomyces* isolates drive developmental reprogramming in *C. neoformans*, related to Figure 2.**

**(A)** Effect of different amounts of tunicamycin on unisexual reproduction in *C. neoformans* strains H99α and KN99**a** during confrontation assay. 0.1, 1.0, 5.0, and 10.0 µg of tunicamycin dissolved in DMSO at different concentrations were spotted onto the disc placed at the center of the plate (replacing the spot of bacterial cells), with 10 µL of DMSO serving as the vehicle control and MS plate only as negative control. Scale bar, 100 µm. **(B)** Light microscopy images showing non-yeast development of *C. neoformans* wild-type strains H99α, KN99**a** and KN99α following confrontation with tunicamycin (TM) and the two tunicamycin-producing *Streptomyces* isolates (WAC1490 and WAC1565) on MS agar. Images were taken after incubation at room temperature in dark for 4 weeks, with those highlighted in red highlighting the production of basidiospore spore chains. Black scale bar, 20 µm; White scale bar, 100 µm. **(C)** Cells of *C. neoformans* H99α were spotted on YPD or MS agar supplemented with tunicamycin at the indicated concentrations and incubated at room temperature in dark for 4 weeks. The top and middle panel show the colony morphologies, and the bottom panel shows light microscopy images of the cells. Tunicamycin of concentration >0.25 μg/mL induced robust non-yeast development, including basidiospore production, although sporulation is impaired when tunicamycin is at concentration above 2.0 μg/mL. Black scale bar, 20 μm; White scale bar, 100 μm. **(D)** Survival of spores and yeast cells exposed to 5 μg/mL TM over 0, 1, 2, and 3 days. Spores were collected from crosses between H99α and KN99**a** on MS medium. Yeast cells were mixtures of the H99α and KN99**a** cells incubated on the same MS medium as the H99α × KN99**a** crosses and were collected at the same time as the basidiospores. **(E)** Cells of *C. neoformans* H99α were spotted on MS agar supplemented with dithiothreitol at the indicated concentrations and incubated at room temperature for 4 weeks in dark. Scale bar, 100 μm. **(F)** Confrontation assays between *C. neoformans* wild-type strain H99α and various antibiotics and antifungals on MS plates in the same way as the confrontation assay of tunicamycin, with one plate per compound. Images show the colony morphology after 4 weeks of incubation at room temperature in the dark. Scale bar, 100 μm. TM, tunicamycin; KM, kanamycin; NEO, Neomycin; CTX, Cefotaxime; Chl, Chloramphenicol; AmB, Amphotericin B; RAPA, Rapamycin; CsA, Cyclosporine; AMA, Antimycin A; CHX, Cycloheximide; ROT, Rotenone; SHAM, Salicylhydroxamic acid.

**Figure S3. Tunicamycin is the causative agent for the induction of unisexual reproduction in *C. neoformans*, related to Figure 2.**

**(A)** Generation of tunicamycin-producing *Streptomyces* strains in the non-tunicamycin-producing host *Streptomyces lividans*. Shown here are LC–MS profiles of culture extracts from *S. lividans* MF1, SBT5, XF8 and XF9 expressing the tunicamycin biosynthetic gene cluster (tunBGC) and their isogenic empty-vector controls. Extracted ion chromatograms were generated for tunicamycin C ($[M+H]^+$ m/z 817.41), tunicamycin V ($[M+H]^+$ m/z 831.42), tunicamycin B ($[M+H]^+$ m/z 845.44), and tunicamycin D ($[M+H]^+$ m/z 859.46) from recombinant *S. lividans* strains carrying the tunBGC or the empty vector control. Diagnostic peaks corresponding to tunicamycins C, V, B, and D are indicated (arrows). Vector controls show no detectable signal at the corresponding retention times. **(B and C)** Confrontation assays between the two tunicamycin-producing *Streptomyces* isolates (WAC1490 and WAC1565), *S. lividans* MF1, SBT5, XF8 and XF9 expressing the tunicamycin biosynthetic gene cluster (tunBGC) and their isogenic empty-vector controls and *C. neoformans* H99α and KN99**a**. The confrontation assays were conducted on MS medium (**B**) and V8 medium (pH=5; **C**). Plates were incubated at room temperature in dark for 4 weeks. Shown here are images of colony morphology at the end of the incubation, with non-yeast developments only observed in *C. neoformans* isolates confronted with tunicamycin-producing natural and engineered *Streptomyces* strains. Scale bar, 100 μm. Vector, isogenic strains carrying empty pRT802; tunBGC, *S. lividans* strains carrying pIJ12003a (pRT802::tunBGC).

**Figure S4. Ploidy analysis of basidiospores produced by unisexual reproduction of *C. neoformans* wild-type haploid (KN99a and H99α) and diploid (BHa/a and BHα/α) strains, related to Figure 3.**

**(A)** FACS analysis of the ploidy of progeny derived from unisexual reproduction of *C. neoformans* wild-type haploid strains H99α or KN99**a** induced by tunicamycin and tunicamycin-producing *Streptomyces* isolates. **(B)** FACS analysis of the ploidy of progeny derived from unisexual reproduction of *C. neoformans* wild-type haploid strains H99α or KN99**a** induced by the engineered S. *lividans* XF8 isolate expressing the tunicamycin biosynthetic gene cluster. **(C)** FACS analysis of the ploidy of progeny derived from unisexual reproduction of *C. neoformans* BH**a**/**a** and BHα/α diploid strains induced by tunicamycin and tunicamycin-producing Streptomyces natural isolates.

**Figure S5. WGS and variant analysis of basidiospores produced from *Streptomyces*- and tunicamycin-induced unisexual reproduction in *C. neoformans* wild-type haploid strains (KN99a and H99α) and diploid strains (BHa/a and BHα/α), related to Figure 3.**

**(A and B)** Read-depth (A) and SNP (B) analyses of progeny derived from unisexual reproduction of *C. neoformans* wild-type haploid strain H99α, using the parental H99α genome as the reference. **(C and D)** Read-depth (C) and SNP (D) analyses of progeny derived from unisexual reproduction of *C. neoformans* wild-type haploid strain KN99**a**, using H99α genome as the reference. The SNP peak on chromosome 5 corresponds to the *MAT* locus, and the SNP peak on chromosome 11 corresponds to a known non-isogenic region between H99α and KN99**a**. **E**, Read depth analyses of progeny dissected from unisexual reproduction of *C. neoformans* BH**a**/**a** or BHα/α diploid strains, using the H99α genome as the reference.

**Figure S6. *Streptomyces*- and tunicamycin-induced unisexual reproduction in *C. neoformans* involved meiotic recombination, related to Figure 3.**
Single-nucleotide polymorphism (SNP) analysis of progeny dissected from unisexual reproduction of *C. neoformans* BH**a**/**a** or BHα/α diploid strains induced by tunicamycin. The plots illustrated the genomic composition of each progeny, with alleles inherited from Bt63**a** shown in blue and those from H99α shown in pink. Each progeny carries a mixture of alleles from both H99α and Bt63**a** across the genome, with frequent crossover events, consistent with their origin through meiosis.

**Figure S7. Analyses of basidiospores confirmed that *Streptomyces* and tunicamycin induced unisexual reproduction in *C. deuterogattii*, related to Figure 7.**
**(A)** FACS analysis of the ploidy of progeny derived from unisexual reproduction of *C. deuterogattii* wild-type strains R265**a** or R265α induced by the natural tunicamycin-producing *Streptomyces* isolates. (**B and C)** Read-depth (B) and SNP (C) analyses of progeny derived from unisexual reproduction of *C. deuterogattii* wild-type haploid strain R265α and R265**a**, using the R265α genome as reference. The SNP peaks on chromosome 9 in progeny dissected from R265**a** correspond to the *MAT* locus.

**Figure S1. Son et al.**

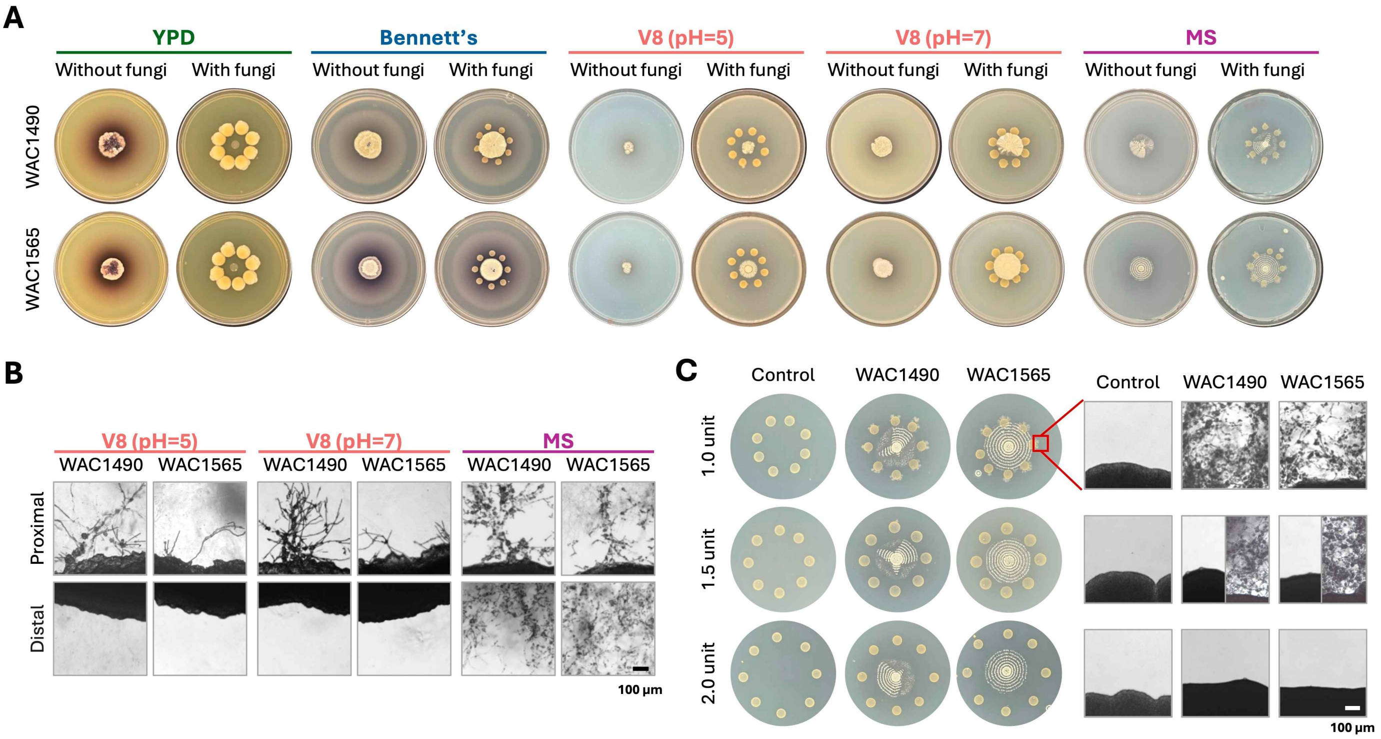

**Figure S2. Son et al.**

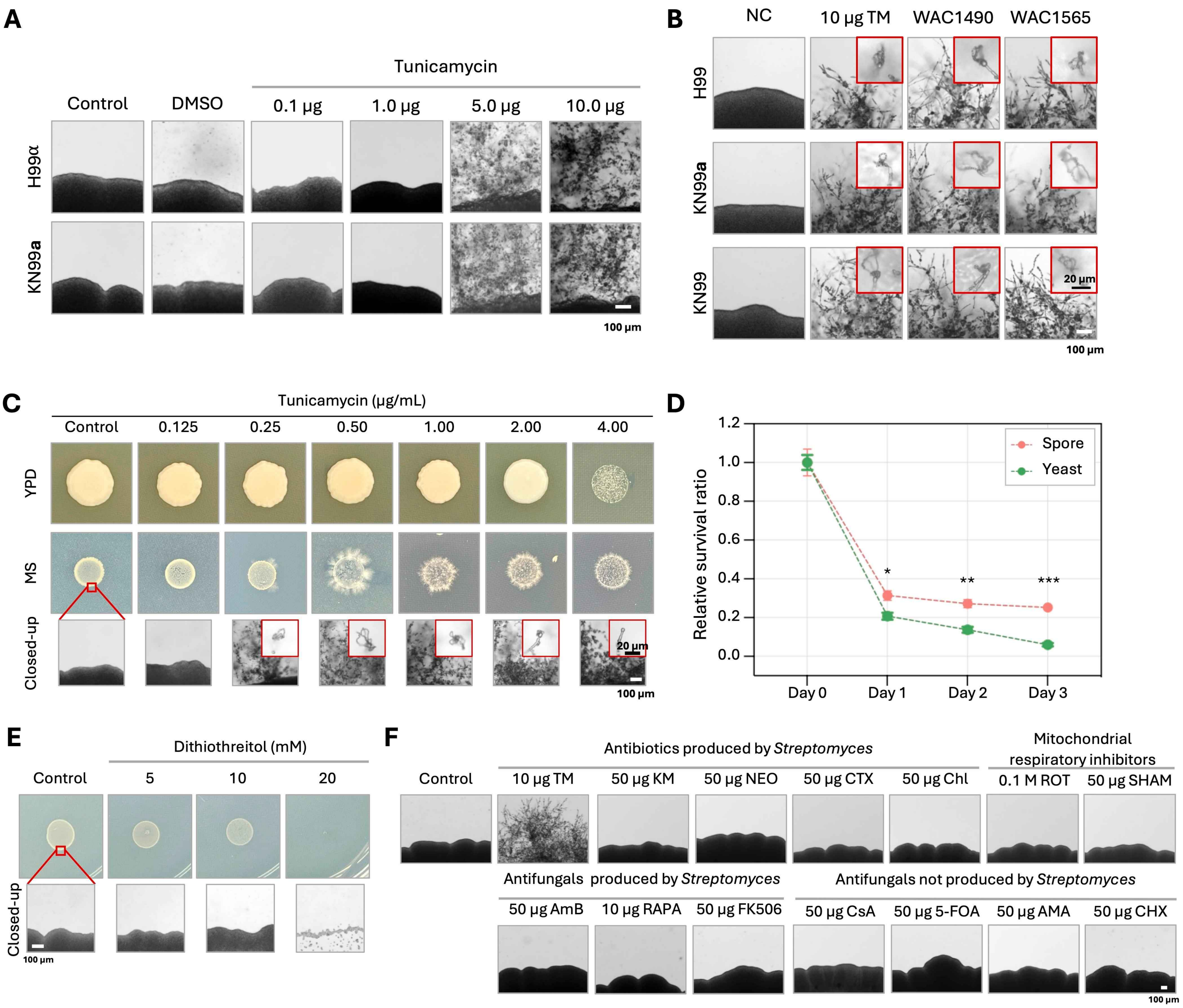

**Figure S3. Son et al.**

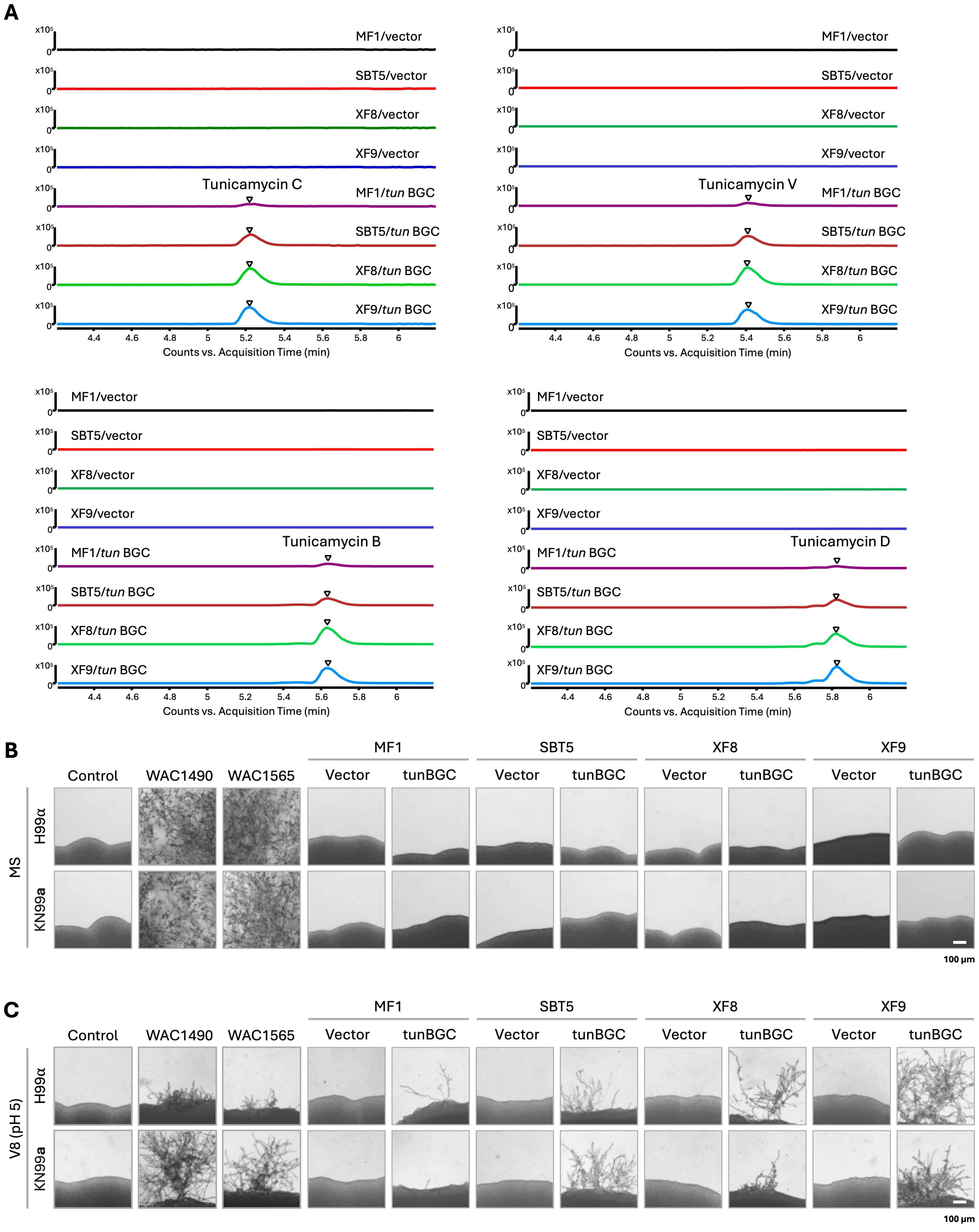

**Figure S4. Son et al.**

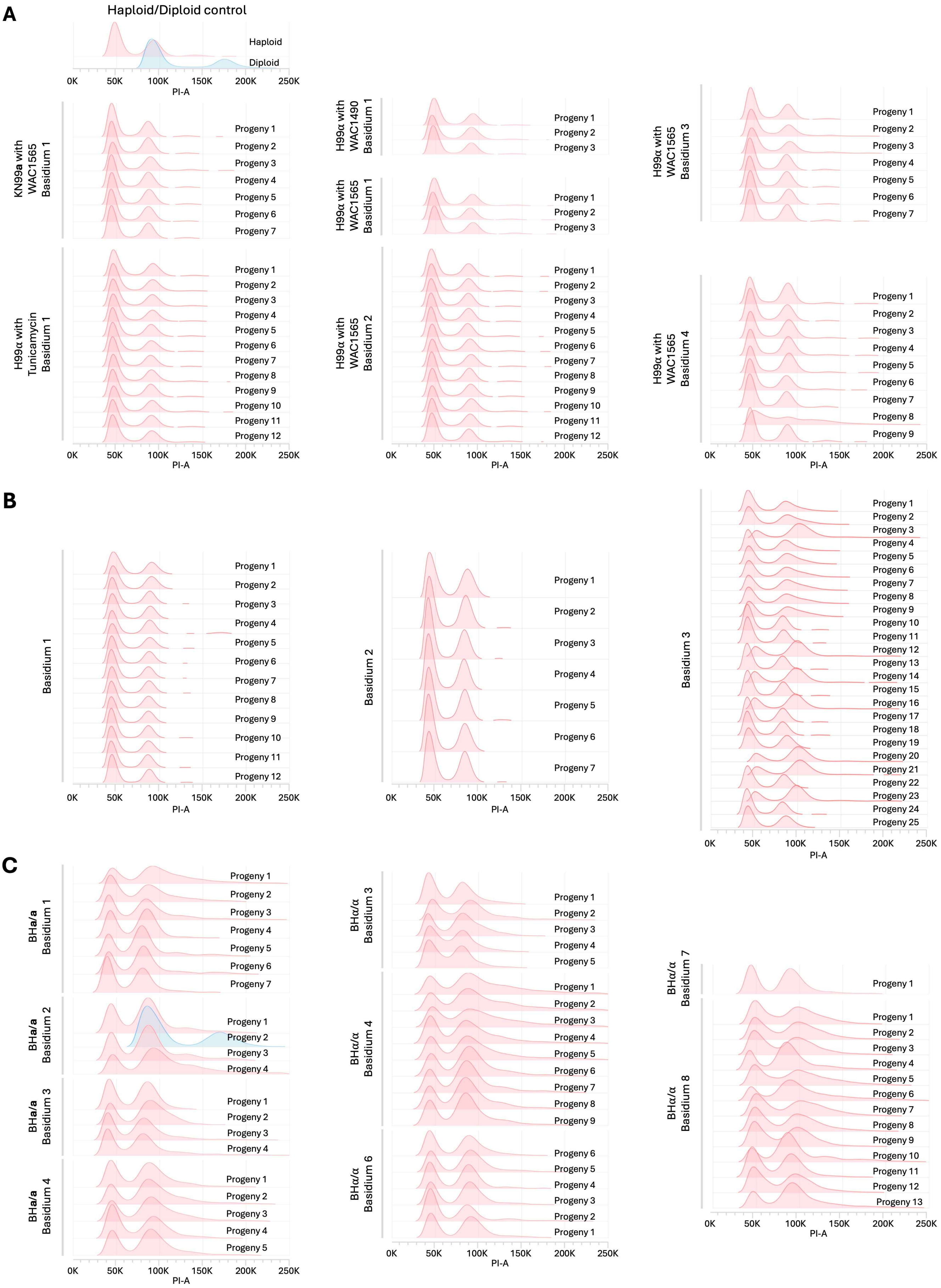

**Figure S5. Son et al.**

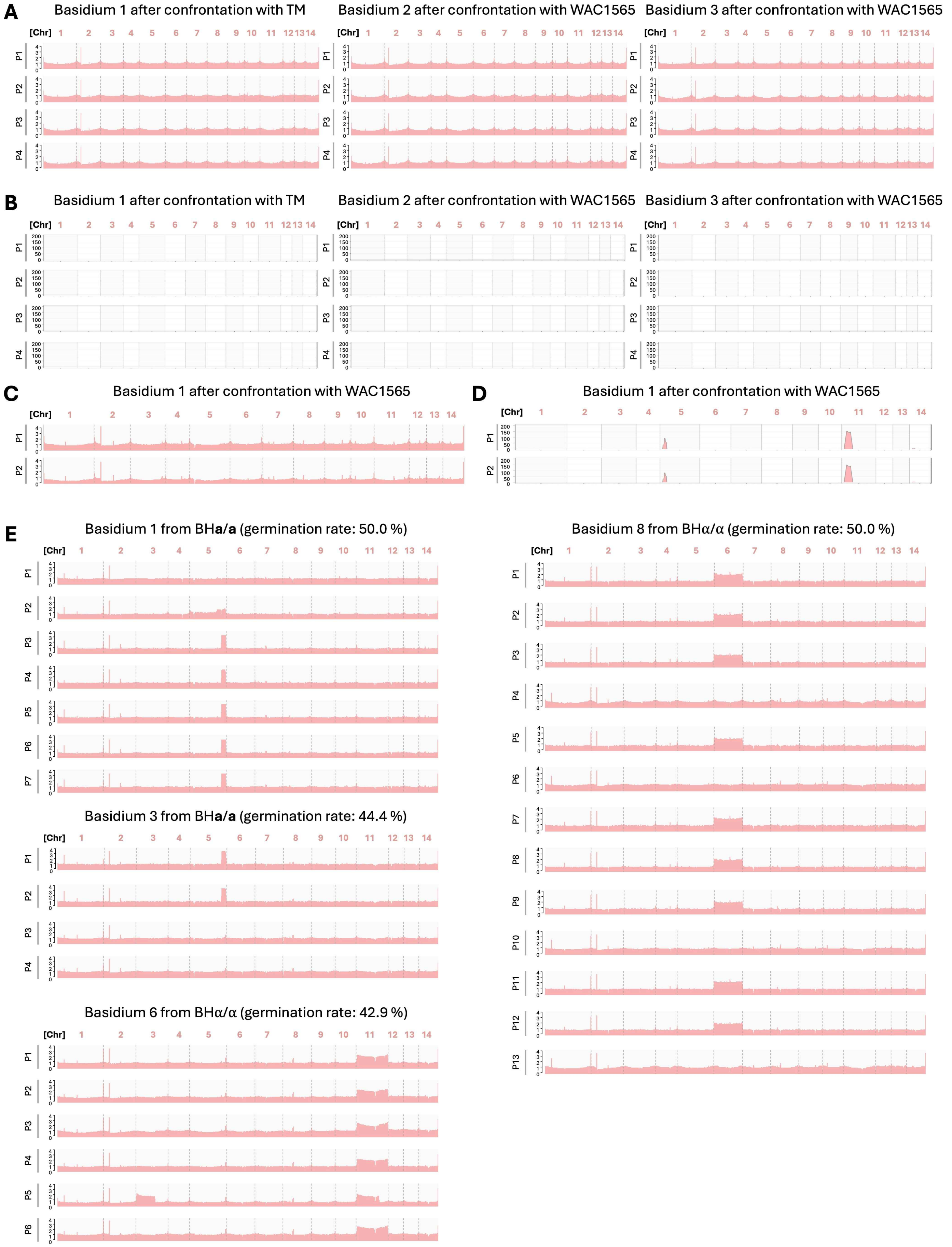

**Figure S6. Son et al.**

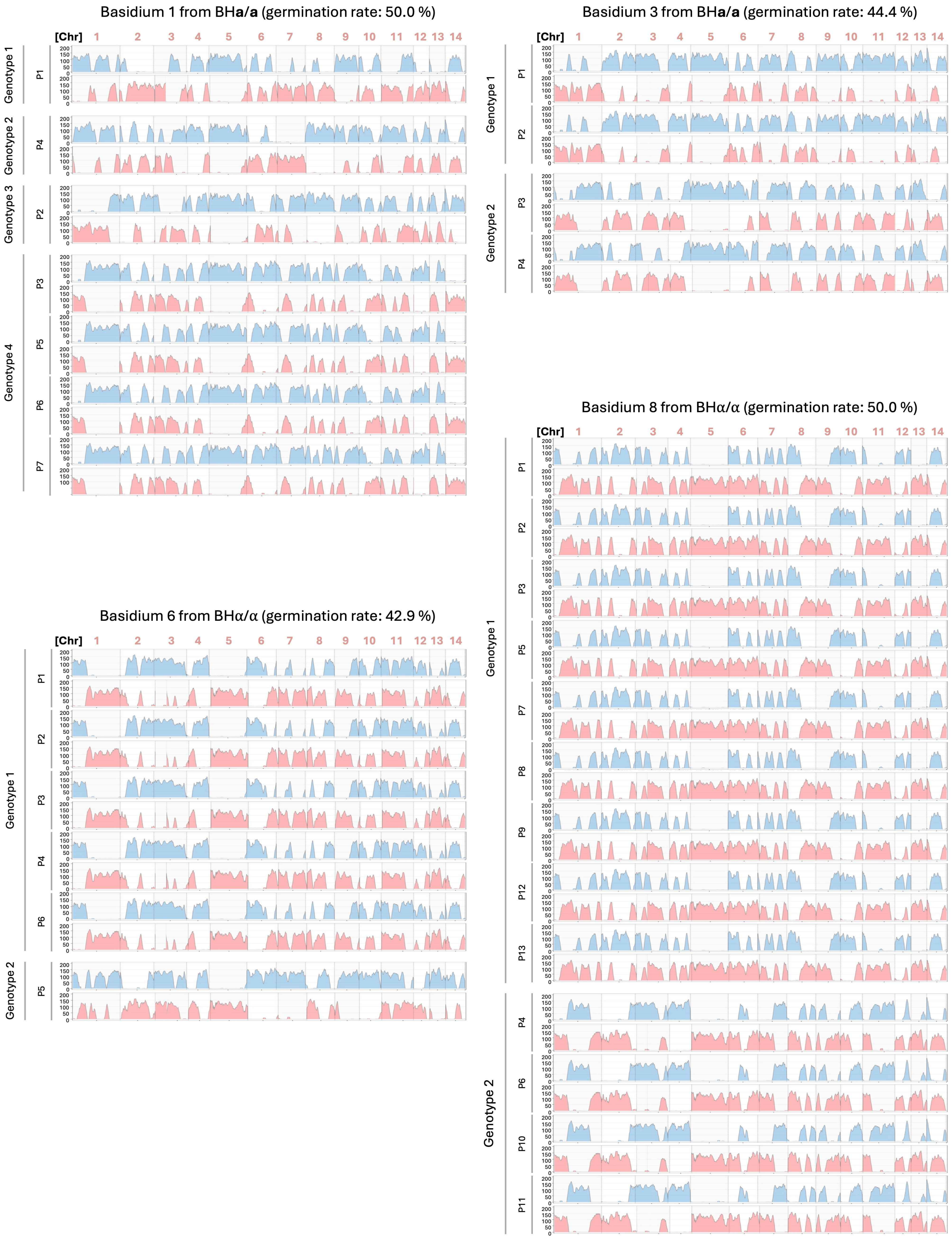

**Figure S7. Son et al.**

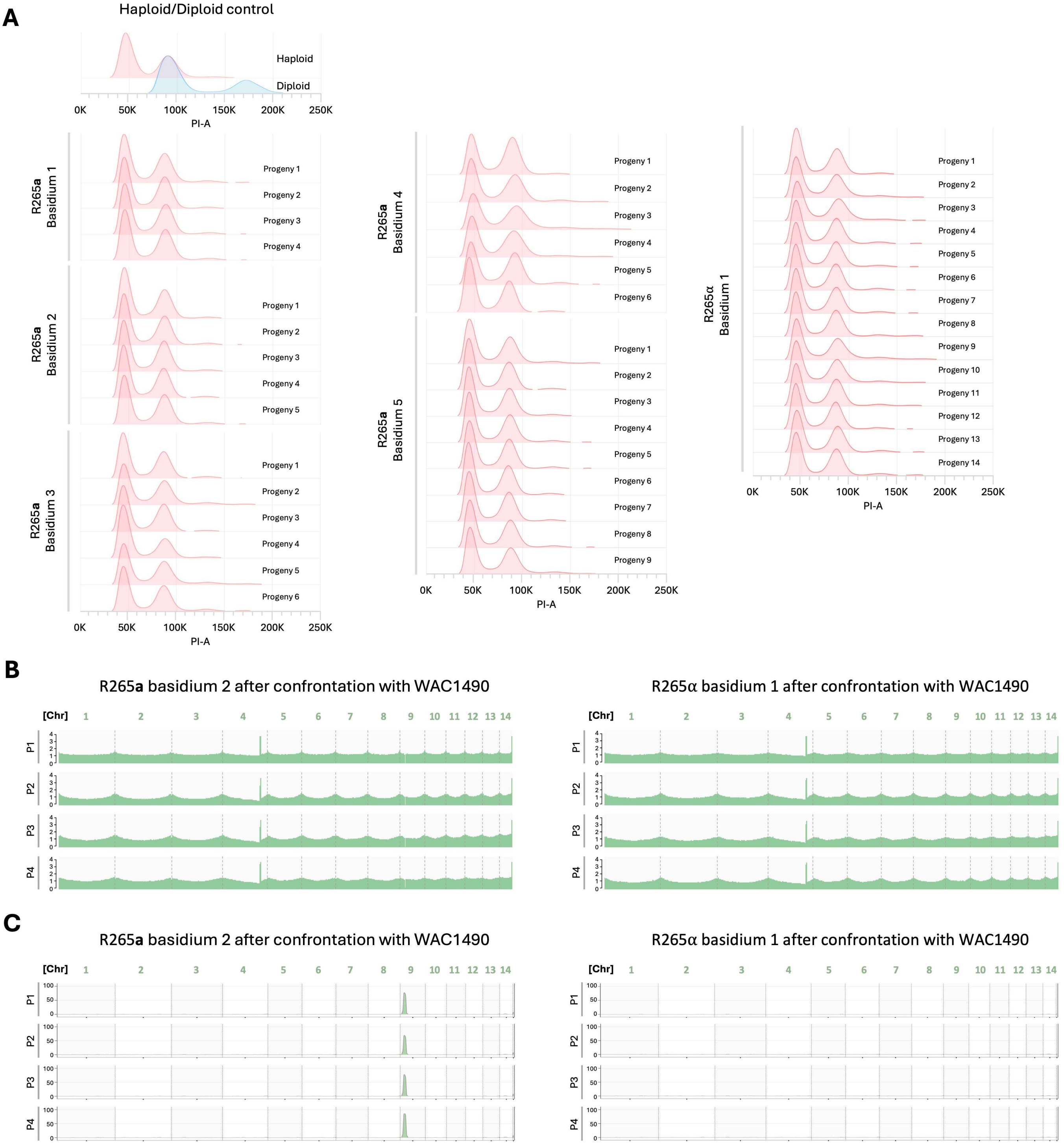

**Table S1. Assessment of bacterial growth on various culture media, related to Figure 1.**

| WAC# | Growth media [1] | | | |
|---|---|---|---|---|
| | Bennett's | YPD | V8 (pH 5) | MS |
| 232 | ○ | ○ | ○ | – |
| 256 | ○ | ○ | ○ | ○ |
| 345 | ○ | ○ | ○ | ○ |
| 409 | ○ | ○ | ○ | ○ |
| 450 | ○ | ○ | ○ | – |
| 466 | ○ | ○ | ○ | ○ |
| 469 | ○ | – | – | – |
| 706 | ○ | ○ | ○ | ○ |
| 759 | ○ | ○ | ○ | – |
| 789 | ○ | ○ | – | – |
| 850 | ○ | ○ | – | – |
| 931 | ○ | ○ | ○ | ○ |
| 1019 | ○ | ○ | ○ | – |
| 1039 | ○ | ○ | – | – |
| 1148 | ○ | ○ | – | – |
| 1490 | ○ | ○ | ○ | ○ |
| 1512 | ○ | ○ | ○ | ○ |
| 1565 | ○ | ○ | ○ | ○ |
| 3260 | ○ | – | – | – |
| 4033 | ○ | ○ | ○ | ○ |
| 4057 | ○ | ○ | ○ | ○ |
| 4093 | ○ | ○ | ○ | ○ |
| 4281 | ○ | ○ | ○ | ○ |
| 4356 | ○ | ○ | ○ | ○ |
| 4488 | ○ | ○ | ○ | ○ |
| 5038 | ○ | ○ | ○ | ○ |
| 6037 | ○ | ○ | ○ | ○ |
| 6068 | ○ | ○ | ○ | ○ |
| 7446 | ○ | ○ | ○ | ○ |
| 7934 | ○ | ○ | ○ | ○ |
| 7987 | ○ | ○ | ○ | ○ |
| 8474 | ○ | ○ | ○ | ○ |
| 8631 | ○ | ○ | ○ | ○ |
| 8659 | ○ | ○ | ○ | – |
| 8921 | ○ | ○ | – | – |
| 9514 | ○ | ○ | ○ | ○ |
| 3273a | ○ | ○ | ○ | – |
| 7751a | ○ | ○ | ○ | ○ |
| 9126a | ○ | – | – | – |

[1] A circle (○) denotes growth on the indicated media, whereas a dash (–) indicates no growth.

**Table S2. Genotypic analysis of unisexual spores derived from *Cryptococcus neoformans* wild-type haploid strains KN99a and H99α and diploid strains BHa/a and BHα/α, related to Figure 3.**

| Fungi | Center | Basidium | Spores | Germinated spores | % Gemination | MAT |
|---|---|---|---|---|---|---|
| KN99**a** | WAC1565 | 1 | 14 | 7 | 50.0 | All **a** |
| H99α | Tunicamycin | 1 | 14 | 12 | 85.7 | All α |
| | WAC1490 | 1 | 8 | 3 | 37.5 | All α |
| | WAC1565 | 1 | 14 | 3 | 21.4 | All α |
| | | 2 | 14 | 12 | 85.7 | All α |
| | | 3 | 8 | 7 | 87.5 | All α |
| | | 4 | 14 | 9 | 64.3 | All α |
| | *S. lividans* XF8 | 1 | 19 | 15 | 78.9 | All α |
| | | 2 | 7 | 7 | 100.0 | All α |
| | | 3 | 28 | 25 | 89.2 | All α |
| BH**a**/**a** | Tunicamycin | 1 | 14 | 7 | 50.0 | All **a** |
| | | 2 | 11 | 4 | 36.4 | All **a** |
| | | 3 | 9 | 4 | 44.4 | All **a** |
| | | 4 | 26 | 5 | 19.2 | All **a** |
| BHα/α | Tunicamycin | 1 | 14 | 0 | 0.0 | All α |
| | | 2 | 14 | 0 | 0.0 | All α |
| | | 3 | 14 | 5 | 35.7 | All α |
| | | 4 | 26 | 9 | 34.6 | All α |
| | | 5 | 14 | 0 | 0.0 | All α |
| | | 6 | 14 | 6 | 42.9 | All α |
| | | 7 | 14 | 1 | 7.1 | All α |
| | | 8 | 26 | 13 | 50.0 | All α |

**Table S3. Summary of genes involved in sexual reproduction and stress response signaling pathways in *C. neoformans*, related to Figure 4.**

| Signaling pathway | Gene name | Description |
|---|---|---|
| Pheromone-responsive MAP kinase (MAPK) | *STE20* | A P21-activated kinase (PAK) required for mating |
| | *STE11* | MAPKKK (Mitogen-Activated Protein Kinase Kinase Kinase) |
| | *STE7* | MAPKK (Mitogen-Activated Protein Kinase Kinase) |
| | *CPK1* | MAPK (Mitogen-Activated Protein Kinase) |
| | *MAT2* | A master regulator of the pheromone-response pathway, A HMG-domain protein required for cell fusion and filamentation |
| cAMP/PKA (Protein Kinase A) | *CAC1* | Adenylyl cyclase |
| | *PKA1* | Protein kinase A |
| Sexual reproduction-related transcription factors | *ZNF2* | Master regulator of controlling yeast-to-hypha dimorphic transition, $C_2H_2$ type zinc finger protein |
| | *SXI1* | Sex-specific homeodomain transcription factor |
| Calcineurin | *CNA1* | Calcineurin A catalytic subunit |
| | *CNB1* | Calcineurin B regulatory subunit |
| | *CRZ1* | Calcineurin-responsive $C_2H_2$ type zinc finger 1 |
| | *PUF4* | A key downstream target of the calcineurin signaling, Pumilio domain-containing RNA-binding protein |
| | *PBP1* | A key downstream target of the calcineurin signaling, Poly(A) binding protein-Binding Protein |
| HOG (High Osmolarity Glycerol) | *SSK1* | An upstream component of the HOG signaling |
| | *SSK2* | MAPKKK (Mitogen-Activated Protein Kinase Kinase Kinase) |
| | *PBS2* | MAPKK (Mitogen-Activated Protein Kinase Kinase) |
| | *HOG1* | MAPK (Mitogen-Activated Protein Kinase) |
| | *ATF1* | A key downstream target of the HOG signaling, bZIP transcription factor |

**Table S4. Commonly regulated differentially expressed genes at early and/or late stages following treatment with tunicamycin or exposure to two tunicamycin-producing *Streptomyces* isolates (WAC1490 and WAC1565), related to Figure 5.**

| Gene ID | Gene name | Description | Titan cell formation | Robust hyphal formation & sporulation |
|---|---|---|---|---|
| *CNAG_00012* | - | D-xylose 1-dehydrogenase (NADP(+), D-xylono-1,5-lactone-forming) | | DOWN |
| *CNAG_00110* | *HPI3* | Formin GTPase-binding domain-containing protein | UP | |
| *CNAG_00126* | - | 2-deoxy-D-gluconate 3-dehydrogenase | | DOWN |
| *CNAG_00162* | *AOX1* | Alternative oxidase, mitochondrial | DOWN | |
| *CNAG_00177* | - | RING-type domain-containing protein | UP | UP |
| *CNAG_00498* | *CDC14* | Protein-tyrosine-phosphatase | UP | |
| *CNAG_00587* | - | Ricin B lectin domain-containing protein | UP | |
| *CNAG_00598* | - | Nicotinamide mononucleotide permease | DOWN | |
| *CNAG_00601* | *CEL1* | AA9 family lytic polysaccharide monooxygenas | UP | UP |
| *CNAG_00726* | - | Major facilitator superfamily (MFS) profile domain-containing protein | | DOWN |
| *CNAG_00729* | - | Helitron helicase-like domain-containing protein | UP | UP |
| *CNAG_00795* | *CFL1* | Protein CPL1-like domain-containing protein | UP | UP |
| *CNAG_00797* | - | Acetyl-coenzyme A synthetase | DOWN | |
| *CNAG_00813* | - | Uncharacterized protein | | UP |
| *CNAG_00814* | - | Transposase domain-containing protein | | DOWN |
| *CNAG_00887* | - | Uncharacterized protein | | UP |
| *CNAG_01081* | - | NAD-dependent epimerase/dehydratase domain-containing protein | UP | |
| *CNAG_01117* | *EF3* | Elongation factor 3 | DOWN | |
| *CNAG_01119* | - | POT family proton-dependent oligopeptide transporter | | UP |
| *CNAG_01122* | - | Calpain catalytic domain-containing protein | UP | |
| *CNAG_01164* | - | Glutamine--fructose-6-phosphate transaminase | UP | |
| *CNAG_01210* | - | Uncharacterized protein | | UP |
| *CNAG_01229* | - | L-lactate dehydrogenase (cytochrome) | | DOWN |
| *CNAG_01257* | - | Aldo-keto reductase | | DOWN |
| *CNAG_01299* | - | Uncharacterized protein | UP | UP |
| *CNAG_01349* | - | MARVEL domain-containing protein | UP | UP |
| *CNAG_01369* | - | Uncharacterized protein | | DOWN |
| *CNAG_01464* | *FHB1* | Nitric oxide dioxygenase | DOWN | |
| *CNAG_01495* | - | Myo-inositol 2-dehydrogenase | | DOWN |
| *CNAG_01512* | - | Carboxylesterase type B domain-containing protein | UP | |
| *CNAG_01572* | *CDC25* | M-phase inducer phosphatase | UP | |
| *CNAG_01653* | *CIG1* | Cytokine inducing-glycoprotein | UP | |
| *CNAG_01668* | - | Peptidase M50B-like-domain-containing protein | UP | UP |
| *CNAG_01794* | - | 2-hydroxyacid dehydrogenase | | DOWN |
| *CNAG_01796* | - | Uncharacterized protein | UP | |
| *CNAG_01803* | - | Uncharacterized protein | | DOWN |
| *CNAG_01806* | - | Uncharacterized protein | UP | UP |
| *CNAG_01816* | - | Uncharacterized protein | UP | |
| *CNAG_01827* | - | VHS domain-containing protein | UP | |
| *CNAG_01995* | - | Extracellular membrane protein CFEM domain-containing protein | | UP |
| *CNAG_02043* | - | VOC domain-containing protein | | DOWN |

| | | | | |
|---|---|---|---|---|
| *CNAG_02053* | - | Uncharacterized protein | DOWN | DOWN |
| *CNAG_02095* | - | Cyclin | UP | |
| *CNAG_02125* | - | Uncharacterized protein | UP | |
| *CNAG_02160* | - | DNA polymerase | UP | |
| *CNAG_02184* | - | Uncharacterized protein | UP | UP |
| *CNAG_02225* | *EXG104* | Glucan 1,3-beta-glucosidase | UP | |
| *CNAG_02288* | - | Solute carrier family 25 | DOWN | |
| *CNAG_02339* | - | Uncharacterized protein | UP | |
| *CNAG_02440* | - | Cation-transporting ATPase | UP | |
| *CNAG_02489* | - | Alcohol dehydrogenase | DOWN | DOWN |
| *CNAG_02499* | - | Uncharacterized protein | UP | |
| *CNAG_02531* | *CPK2* | Calcium-dependent protein kinase 2 | UP | |
| *CNAG_02562* | - | Acyl-CoA dehydrogenase | | DOWN |
| *CNAG_02694* | - | Ricin B lectin domain-containing protein | UP | |
| *CNAG_02701* | - | BAG domain-containing protein | UP | UP |
| *CNAG_02735* | - | Uncharacterized protein | | DOWN |
| *CNAG_02777* | *PHO84* | Phosphate:H symporter | DOWN | |
| *CNAG_02839* | - | FAD/NAD(P)-binding domain-containing protein | UP | |
| *CNAG_02863* | - | Formate dehydrogenase | DOWN | DOWN |
| *CNAG_02865* | - | WSC domain-containing protein | UP | |
| *CNAG_02899* | - | Uncharacterized protein | DOWN | |
| *CNAG_03021* | - | Uncharacterized protein | UP | UP |
| *CNAG_03037* | - | Uncharacterized protein | UP | |
| *CNAG_03047* | - | DUF300-domain-containing protein | UP | |
| *CNAG_03068* | - | Methyltransferase domain-containing protein | UP | |
| *CNAG_03107* | - | Uncharacterized protein | UP | UP |
| *CNAG_03125* | - | Hyaluronan-mediated motility receptor C-terminal domain-containing protein | UP | |
| *CNAG_03176* | - | Endoplasmic oxidoreductin 1 | UP | |
| *CNAG_03223* | - | Yeast cell wall synthesis Kre9/Knh1-like N-terminal domain-containing protein | UP | |
| *CNAG_03224* | - | Uncharacterized protein | UP | UP |
| *CNAG_03272* | - | 3-oxoacyl-[acyl-carrier-protein] reductase | | UP |
| *CNAG_03277* | - | Inositol oxygenase | | DOWN |
| *CNAG_03292* | - | RRM domain-containing protein | DOWN | |
| *CNAG_03316* | *RDI1* | Rho GDP-dissociation inhibitor | UP | |
| *CNAG_03384* | - | Uncharacterized protein | | UP |
| *CNAG_03443* | - | Uncharacterized protein | UP | |
| *CNAG_03453* | *KIP1* | Kinesin family member 11 | UP | |
| *CNAG_03465* | *LAC1* | Laccase-1 | | DOWN |
| *CNAG_03539* | - | SprT-like domain-containing protein | UP | |
| *CNAG_03617* | *CLP1* | Clampless protein 1 | UP | UP |
| *CNAG_03630* | - | Uncharacterized protein | | UP |
| *CNAG_03683* | - | Short-chain dehydrogenase/reductase SDR | | DOWN |
| *CNAG_03714* | - | Microtubule motor protein | UP | |
| *CNAG_03741* | *FZC31* | Zn(2)-C6 fungal-type domain-containing protein | UP | |
| *CNAG_03744* | - | Uncharacterized protein | UP | |
| *CNAG_03755* | - | Uncharacterized protein | UP | |

| | | | | |
|---|---|---|---|---|
| *CNAG_03833* | - | BTB domain-containing protein | UP | |
| *CNAG_03980* | - | BTB domain-containing protein | UP | UP |
| *CNAG_04007* | - | Septin ring protein | UP | UP |
| *CNAG_04015* | - | Amino acid transporter | | DOWN |
| *CNAG_04018* | - | Subtelomeric hrmA-associated cluster protein | UP | |
| *CNAG_04085* | - | Oxidoreductase | UP | |
| *CNAG_04139* | - | Uncharacterized protein | | DOWN |
| *CNAG_04210* | - | MFS transporter, SP family, general alpha glucoside:H symporter | | DOWN |
| *CNAG_04245* | *CHI22* | Chitinase | UP | |
| *CNAG_04342* | - | Uncharacterized protein | UP | |
| *CNAG_04344* | - | Galactonate dehydratase | | DOWN |
| *CNAG_04346* | - | Dihydrodipicolinate synthase | | DOWN |
| *CNAG_04357* | - | Yeast cell wall synthesis Kre9/Knh1-like N-terminal domain-containing protein | UP | UP |
| *CNAG_04467* | - | Succinate-semialdehyde dehydrogenase | UP | |
| *CNAG_04548* | - | DNA 3'-5' helicase (EC 5624) | | UP |
| *CNAG_04677* | - | Xylitol dehydrogenase | | DOWN |
| *CNAG_04696* | - | DNA clamp loader | UP | |
| *CNAG_04707* | - | CFEM domain-containing protein | UP | |
| *CNAG_04736* | - | Uncharacterized protein | UP | |
| *CNAG_04759* | - | Uncharacterized protein | | UP |
| *CNAG_04857* | - | Uncharacterized protein | UP | |
| *CNAG_04861* | - | Transglycosylase SLT domain-containing protein | UP | UP |
| *CNAG_04891* | - | Ricin B lectin domain-containing protein | UP | |
| *CNAG_04892* | - | F-box domain-containing protein | UP | |
| *CNAG_04921* | - | Uncharacterized protein | | DOWN |
| *CNAG_04975* | - | Uncharacterized protein | | UP |
| *CNAG_05010* | *ZFC7* | C2H2-type domain-containing protein | | UP |
| *CNAG_05032* | - | Uncharacterized protein | UP | |
| *CNAG_05072* | - | Endopeptidase | UP | UP |
| *CNAG_05075* | - | Phosphate transporter | UP | |
| *CNAG_05099* | - | Uncharacterized protein | UP | |
| *CNAG_05264* | - | Alpha-amylase AmyA | UP | |
| *CNAG_05265* | - | Uncharacterized protein | UP | |
| *CNAG_05303* | *ICL1* | Isocitrate lyase | DOWN | |
| *CNAG_05305* | - | Uncharacterized protein | UP | UP |
| *CNAG_05326* | - | Transcription factor domain-containing protein | DOWN | |
| *CNAG_05330* | - | MFS transporter, SP family, general alpha glucoside:H symporter | | DOWN |
| *CNAG_05340* | - | Monosaccharide transporter | | DOWN |
| *CNAG_05351* | *PMM1* | Phosphomannomutase | UP | |
| *CNAG_05352* | - | Acetyl-CoA transporter | UP | |
| *CNAG_05381* | *ITR3C* | Myo-inositol transporter 3C | | DOWN |
| *CNAG_05412* | - | Uncharacterized protein | | UP |
| *CNAG_05448* | - | Uncharacterized protein | UP | |
| *CNAG_05458* | - | Endo-1,3(4)-beta-glucanase | UP | |
| *CNAG_05461* | - | L-lactate dehydrogenase | | DOWN |
| *CNAG_05514* | - | Uncharacterized protein | UP | UP |

| | | | | |
|---|---|---|---|---|
| *CNAG_05565* | - | Uncharacterized protein | UP | |
| *CNAG_05576* | - | Uncharacterized protein | UP | |
| *CNAG_05592* | *ECA1* | P-type Ca(2+) transporter | UP | |
| *CNAG_05602* | *PUT2* | Multifunctional fusion protein | UP | |
| *CNAG_05637* | - | Spo12 | UP | |
| *CNAG_05653* | *MLS1* | Malate synthase | DOWN | |
| *CNAG_05668* | - | Uncharacterized protein | UP | |
| *CNAG_05678* | - | Membrane protein | DOWN | DOWN |
| *CNAG_05724* | - | Uncharacterized protein | UP | |
| *CNAG_05729* | - | Uncharacterized protein | UP | UP |
| *CNAG_05741* | - | Thioesterase | | DOWN |
| *CNAG_05760* | - | Endoplasmic reticulum protein | | DOWN |
| *CNAG_05785* | *STB4* | Zn(2)-C6 fungal-type domain-containing protein | | DOWN |
| *CNAG_05885* | - | Uncharacterized protein | UP | UP |
| *CNAG_05914* | - | MFS transporter, SP family, general alpha glucoside:H symporter | DOWN | |
| *CNAG_05972* | - | Uncharacterized protein | UP | |
| *CNAG_06082* | - | Delayed-type hypersensitivity antigen | UP | |
| *CNAG_06239* | - | Uncharacterized protein | | UP |
| *CNAG_06240* | - | Protein disulfide-isomerase | UP | |
| *CNAG_06297* | - | Uncharacterized protein | DOWN | |
| *CNAG_06298* | - | Uncharacterized protein | DOWN | |
| *CNAG_06312* | - | Uncharacterized protein | UP | |
| *CNAG_06336* | - | Glucan 1,3-beta-glucosidase | UP | |
| *CNAG_06443* | *KAR2* | Heat shock 70 kDa protein C | UP | |
| *CNAG_06527* | - | MFS transporter, SP family, general alpha glucoside:H symporter | | DOWN |
| *CNAG_06529* | - | D-xylose 1-dehydrogenase (NADP(+), D-xylono-1,5-lactone-forming) | | DOWN |
| *CNAG_06531* | - | Zn(2)-C6 fungal-type domain-containing protein | | DOWN |
| *CNAG_06551* | - | Carnitine O-acetyltransferase | DOWN | DOWN |
| *CNAG_06561* | - | Allantoate transporter | DOWN | DOWN |
| *CNAG_06583* | - | Uncharacterized protein | DOWN | |
| *CNAG_06590* | - | Uncharacterized protein | | DOWN |
| *CNAG_06658* | - | Rhomboid-type serine protease | UP | |
| *CNAG_06659* | *HEX1* | Beta-hexosaminidase | UP | |
| *CNAG_06662* | - | Uncharacterized protein | UP | UP |
| *CNAG_06715* | - | Uncharacterized protein | UP | UP |
| *CNAG_06718* | - | Uncharacterized protein | | UP |
| *CNAG_06795* | - | Yeast cell wall synthesis Kre9/Knh1-like N-terminal domain-containing protein | UP | |
| *CNAG_06814* | *SXI1alpha* | KN homeodomain domain-containing protein | UP | |
| *CNAG_06833* | - | CxC1-like cysteine cluster | | UP |
| *CNAG_06902* | *CPK1* | Mitogen-activated protein kinase | UP | |
| *CNAG_06949* | - | HMG box domain-containing protein | | DOWN |
| *CNAG_07399* | - | Uncharacterized protein | UP | UP |
| *CNAG_07406* | - | Pheromone alpha | UP | UP |
| *CNAG_07407* | *MFalpha3* | Fungal mating-type pheromone | UP | UP |
| *CNAG_07589* | - | Xylitol dehydrogenase | | DOWN |
| *CNAG_07592* | - | Uncharacterized protein | UP | UP |

| | | | | |
|---|---|---|---|---|
| *CNAG_07648* | - | Uncharacterized protein | UP | |
| *CNAG_07736* | - | Glucan endo-1,3-alpha-glucosidase agn1 | | DOWN |
| *CNAG_07747* | - | Acyl-coenzyme A oxidase | | DOWN |
| *CNAG_07756* | - | Cell division control protein | UP | |
| *CNAG_07795* | - | Uncharacterized protein | UP | UP |
| *CNAG_07832* | - | Uncharacterized protein | DOWN | |
| *CNAG_07842* | - | V-SNARE coiled-coil homology domain-containing protein | UP | |
| *CNAG_07845* | - | Uncharacterized protein | DOWN | |
| *CNAG_07850* | - | Uncharacterized protein | | UP |
| *CNAG_07916* | - | Uncharacterized protein | | UP |
| *CNAG_07943* | - | Store-operated calcium entry-associated regulatory factor | UP | |
| *CNAG_07981* | - | Uncharacterized protein | UP | UP |
| *CNAG_08015* | - | Cysteine-rich transmembrane CYSTM domain-containing protein | UP | |

**Table S5. Top 20 differentially up-expressed genes after titan cell formation, related to Figure 5.**

| Gene ID | Gene name | $Log_2$(Fold change) | | | Description |
|---|---|---|---|---|---|
| | | vs TM | vs WAC1490 | vs WAC1590 | |
| *CNAG_00795* | *CFL1* | 9.43 | 9.64 | 9.59 | Protein CPL1-like domain-containing protein |
| *CNAG_04007* | - | 7.09 | 5.97 | 6.02 | Septin ring protein |
| *CNAG_05448* | - | 6.30 | 4.80 | 6.10 | Uncharacterized protein |
| *CNAG_06312* | - | 6.16 | 6.50 | 6.26 | Uncharacterized protein |
| *CNAG_03741* | *FZC31* | 6.07 | 5.01 | 5.30 | Zn(2)-C6 fungal-type domain-containing protein |
| *CNAG_05729* | - | 5.96 | 4.50 | 4.97 | Uncharacterized protein |
| *CNAG_05305* | - | 5.93 | 3.70 | 3.69 | Uncharacterized protein |
| *CNAG_07406* | - | 5.72 | 6.23 | 6.00 | Pheromone alpha |
| *CNAG_07399* | - | 5.45 | 4.22 | 4.38 | Uncharacterized protein |
| *CNAG_07407* | *MFalpha3* | 5.22 | 5.51 | 5.30 | Fungal mating-type pheromone |
| *CNAG_01806* | - | 4.89 | 4.38 | 3.85 | Uncharacterized protein |
| *CNAG_03833* | - | 4.72 | 3.27 | 3.45 | BTB domain-containing protein |
| *CNAG_02339* | - | 4.69 | 3.40 | 3.21 | Uncharacterized protein |
| *CNAG_02499* | - | 4.44 | 3.99 | 4.16 | Uncharacterized protein |
| *CNAG_03107* | - | 4.35 | 2.52 | 2.73 | Uncharacterized protein |
| *CNAG_04861* | - | 4.25 | 3.23 | 3.32 | Transglycosylase SLT domain-containing protein |
| *CNAG_01653* | *CIG1* | 4.03 | 3.35 | 3.56 | Cytokine inducing-glycoprotein |
| *CNAG_01816* | - | 4.00 | 3.71 | 3.69 | Uncharacterized protein |
| *CNAG_07943* | - | 3.89 | 3.56 | 3.24 | Store-operated calcium entry-associated regulatory factor |
| *CNAG_06082* | - | 3.84 | 3.18 | 2.77 | Delayed-type hypersensitivity antigen |

**Table S6. Genotypic analysis of unisexual spores produced from *Cryptococcus deuterogattii* wild-type R265a or R265α, related to Figure 7**

| Fungi | Center | Basidium | Spores | Germinated spores | % Gemination | MAT |
|---|---|---|---|---|---|---|
| R265α | WAC1490 | 1 | 14 | 14 | 100.0 | All SXI1α |
| R265**a** | WAC1490 | 1 | 8 | 4 | 50.0 | All SXI2**a** |
| | | 2 | 6 | 5 | 83.3 | All SXI2**a** |
| | | 3 | 14 | 6 | 42.9 | All SXI2**a** |
| | | 4 | 13 | 6 | 46.2 | All SXI2**a** |
| | | 5 | 14 | 9 | 64.3 | All SXI2**a** |

### Table S7. *Streptomyces* strains used for confrontation assay.

| WAC# | Tunicamycin biosynthetic gene cluster[†] | Closest genus | Closest species | Source |
|---|---|---|---|---|
| 232 | – | *Streptomyces* | *lunaelactis* | Canada |
| 256 | – | *Streptomyces* | *anulatus* | Canada |
| 345 | – | *Streptomyces* | *violaceoruber* | Canada |
| 409 | – | *Streptomyces* | *anulatus* | Canada |
| 450 | – | *Streptomyces* | *erythrochromogenes* | Canada |
| 466 | – | *Streptomyces* | *murinus* | Canada |
| 706 | – | *Kitasatospora* | *sp001625365* | Canada |
| 759 | – | *Streptomyces* | *sp940670765* | Canada |
| 931 | – | *Streptomyces* | *sp000719575* | Canada |
| 1019 | – | *Streptomyces* | *sp040652845* | Canada |
| 1490 | ○ | *Streptomyces* | *cyaneofuscatus* | Canada |
| 1512 | – | *Streptomyces* | *globisporus* | Canada |
| 1565 | ○ | *Streptomyces* | *cyaneofuscatus* | Canada |
| 3273a | | *Streptomyces* | *olivaceus* | Canada |
| 4033 | – | *Streptomyces* | *anulatus* | Canada |
| 4057 | – | *Streptomyces* | *sp.* | USA |
| 4093 | – | *Streptomyces* | *cyaneofuscatus* | USA |
| 4281 | – | *Streptomyces* | *niveus* | Canada |
| 4356 | – | *Streptomyces* | *werraensis* | Canada |
| 4488 | – | *Streptomyces* | *sp040609055* | Canada |
| 5038 | – | *Streptomyces* | *hygroscopicus* | Canada |
| 6037 | – | *Streptomyces* | *sanglieri* | Canada |
| 6068 | – | *Streptomyces* | *sp001905125* | Canada |
| 7446 | – | *Streptomyces* | *cyaneofuscatus* | Canada |
| 7751a | – | *Streptomyces* | *microflavus* | Canada |
| 7934 | – | *Streptomyces* | *microflavus* | Canada |
| 7987 | – | *Streptomyces* | *microflavus* | Canada |
| 8474 | – | *Streptomyces* | *anulatus* | Canada |
| 8631 | – | *Streptomyces* | *sp001905125* | Canada |
| 8659 | – | *Streptomyces* | *sp040607125* | Canada |
| 9514 | – | *Streptomyces* | *sp001905125* | Canada |

[†] A circle (○) indicates presence, whereas a dash (–) indicates absence.

### Table S8. *Cryptococcus* strains analyzed in this study.

| Species | Strain name | Relevant genotype | Source/ references |
|---|---|---|---|
| *Cryptococcus var. grubii* (VNI) | H99α | *MAT*α wild type | [1] |
| *Cryptococcus var. grubii* (VNI) | KN99**a** | *MAT***a** wild type | [2] |
| *Cryptococcus var. grubii* (VNI) | KN99α | *MAT*α wild type | [2] |
| *Cryptococcus var. grubii* (VNI) | BH**a**/**a** | BH**a**/**a** heterozygous diploid | This study |
| *Cryptococcus var. grubii* (VNI) | BHα/α | BHα/α heterozygous diploid | This study |
| *Cryptococcus var. grubii* (VNI) | SSL791 | *MAT*α pDMC1-GFP-tH3 | This study |
| *Cryptococcus var. grubii* (VNI) | YSB552 | *MAT*α *ire1*△::*NAT*-STM#224 | [3] |
| *Cryptococcus var. grubii* (VNI) | YSB554 | *MAT*α *ire1*△::*NAT*-STM#224 | [3] |
| *Cryptococcus var. grubii* (VNI) | YSB723 | *MAT*α *hxl1*△::*NAT*-STM195 | [3] |
| *Cryptococcus var. grubii* (VNI) | CK0191 | *MAT*α *hxl1*△::*NAT* | [4] |
| *Cryptococcus var. grubii* (VNI) | SSI867 | *MAT*α *ric8*△::*NAT* | This study |
| *Cryptococcus var. grubii* (VNI) | SSK080 | *MAT*α *ric8*△::*NAT*; *ste3*△::*NEO* | This study |
| *Cryptococcus var. grubii* (VNI) | CDX6 | *MAT*α *gpr4*△::*NEO* | [5] |
| *Cryptococcus var. grubii* (VNI) | CDX7 | *MAT*α *gpr4*△::*NAT209* | [5] |
| *Cryptococcus var. grubii* (VNI) | YSB84 | *MAT*α *gpa1*△::*NAT-STM#5* | [6] |
| *Cryptococcus var. grubii* (VNI) | YSB85 | *MAT*α *gpa1*△::*NAT-STM#5* | [6] |
| *Cryptococcus var. grubii* (VNI) | YSB25 | *MAT*α *gpa2*△::*NEO* | [7] |
| *Cryptococcus var. grubii* (VNI) | YSB26 | *MAT*α *gpa2*△::*NEO* | [7] |
| *Cryptococcus var. grubii* (VNI) | JF67 | *MAT*α *gpa3*△::*NAT3.3* | [8] |
| *Cryptococcus var. grubii* (VNI) | CK0492 | *MAT*α *gpa3*△::NAT | [4] |
| *Cryptococcus var. grubii* (VNI) | YSB48 | *MAT*α *gpb1*△::*NAT-STM#146* | [6] |
| *Cryptococcus var. grubii* (VNI) | YSB50 | *MAT*α *gpb1*△::*NAT-STM#146* | [6] |
| *Cryptococcus var. grubii* (VNI) | PPW94 | *MAT*α *ste20*△::*URA5* | [9] |
| *Cryptococcus var. grubii* (VNI) | PPW95 | *MAT*α *ste20*△::*URA5* | [9] |
| *Cryptococcus var. grubii* (VNI) | YSB313 | *MAT*α *ste11*△::*NAT-STM#242* | [10] |
| *Cryptococcus var. grubii* (VNI) | YSB314 | *MAT*α *ste11*△::*NAT-STM#242* | [10] |
| *Cryptococcus var. grubii* (VNI) | YSB342 | *MAT*α *ste7*△::*NAT-STM#225* | [11] |
| *Cryptococcus var. grubii* (VNI) | YSB343 | *MAT*α *ste7*△::*NAT-STM#225* | [11] |
| *Cryptococcus var. grubii* (VNI) | YSB127 | *MAT*α *cpk1*△::*NAT-STM#184* | [11] |
| *Cryptococcus var. grubii* (VNI) | YSB128 | *MAT*α *cpk1*△::*NAT-STM#184* | [11] |
| *Cryptococcus var. grubii* (VNI) | XL1598 | *MAT*α *mat2*△::*NEO* | [12] |
| *Cryptococcus var. grubii* (VNI) | CK0146 | *MAT*α *mat2*△::*NAT* | [4] |
| *Cryptococcus var. grubii* (VNI) | YSB42 | *MAT*α *cac1*△::*NAT-STM#43* | [13] |
| *Cryptococcus var. grubii* (VNI) | YSB5650 | *MAT*α *cac1*△::*NAT* | [13] |
| *Cryptococcus var. grubii* (VNI) | YSB188 | *MAT*α *pka1*△::*NAT-STM#191* | [6] |
| *Cryptococcus var. grubii* (VNI) | YSB189 | *MAT*α *pka1*△::*NAT-STM#191* | [6] |
| *Cryptococcus var. grubii* (VNI) | XL1601 | *MAT*α *znf2*△::*NEO* | [12] |
| *Cryptococcus var. grubii* (VNI) | CK0135 | *MAT*α *znf2*△::*NAT* | [4] |
| *Cryptococcus var. grubii* (VNI) | YSB1391 | *MAT*α *sxi1*△::*NAT*-STM#208 | [14] |
| *Cryptococcus var. grubii* (VNI) | YSB1392 | *MAT*α *sxi1*△::*NAT*-STM#208 | [14] |
| *Cryptococcus var. grubii* (VNI) | YFF1 | *MAT*α *dmc1*△::*NAT* | This study |
| *Cryptococcus var. grubii* (VNI) | YFF2 | *MAT*α *dmc1*△::*NAT* | This study |
| *Cryptococcus var. grubii* (VNI) | ZB301 | *MAT*α *spo11*△::*NAT* | [15] |
| *Cryptococcus var. grubii* (VNI) | SSM615 | *MAT*α *spo11*△::*NAT* | This study |
| *Cryptococcus var. grubii* (VNI) | HP242 | *MAT*α *cna1*△::*NEO* | [16] |
| *Cryptococcus var. grubii* (VNI) | YSB4067 | *MAT*α *cna1*△::*NAT-STM#290* | [13] |

| | | | |
|---|---|---|---|
| *Cryptococcus var. grubii* (VNI) | KK2 | *MAT*α *cnb1*△::*NAT* | 17 |
| *Cryptococcus var. grubii* (VNI) | KK18 | *MAT*α *cnb1*△::*NAT* | 17 |
| *Cryptococcus var. grubii* (VNI) | HP235 | *MAT*α *crz1*△::*NEO* | 16 |
| *Cryptococcus var. grubii* (VNI) | YSB1263 | *MAT*α *crz1*△::*NAT-STM#231* | 14 |
| *Cryptococcus var. grubii* (VNI) | HP17 | *MAT*α *puf4*△::*NEO* | 16 |
| *Cryptococcus var. grubii* (VNI) | HP19 | *MAT*α *puf4*△::*NEO* | 16 |
| *Cryptococcus var. grubii* (VNI) | HP5 | *MAT*α *pbp1*△::*NEO* | 16 |
| *Cryptococcus var. grubii* (VNI) | HP7 | *MAT*α *pbp1*△::*NEO* | 16 |
| *Cryptococcus var. grubii* (VNI) | YSB278 | *MAT*α *tco1*△::*NAT-STM#102* | 18 |
| *Cryptococcus var. grubii* (VNI) | YSB279 | *MAT*α *tco1*△::*NAT-STM#102* | 18 |
| *Cryptococcus var. grubii* (VNI) | YSB281 | *MAT*α *tco2*△::*NAT-STM#116* | 18 |
| *Cryptococcus var. grubii* (VNI) | YSB282 | *MAT*α *tco2*△::*NAT-STM#116* | 18 |
| *Cryptococcus var. grubii* (VNI) | YSB261 | *MAT*α *ssk1*△::*NAT-STM#205* | 18 |
| *Cryptococcus var. grubii* (VNI) | YSB262 | *MAT*α *ssk1*△::*NAT-STM#205* | 18 |
| *Cryptococcus var. grubii* (VNI) | YSB264 | *MAT*α *ssk2*△::*NAT-STM#210* | 10 |
| *Cryptococcus var. grubii* (VNI) | YSB265 | *MAT*α *ssk2*△::*NAT-STM#210* | 10 |
| *Cryptococcus var. grubii* (VNI) | YSB123 | *MAT*α *pbs2*△::*NAT-STM#213* | 19 |
| *Cryptococcus var. grubii* (VNI) | YSB124 | *MAT*α *pbs2*△::*NAT-STM#213* | 19 |
| *Cryptococcus var. grubii* (VNI) | YSB64 | *MAT*α *hog1*△::*NAT-STM#177* | 19 |
| *Cryptococcus var. grubii* (VNI) | CK0467 | *MAT*α *hog1*△::*NAT* | 4 |
| *Cryptococcus var. grubii* (VNI) | YSB676 | *MAT*α *atf1*△::*NAT-STM#220* | 20 |
| *Cryptococcus var. grubii* (VNI) | YSB678 | *MAT*α *atf1*△::*NAT-STM#220* | 20 |
| *Cryptococcus var. grubii* (VNI) | CK3320 | *MAT*α *CNAG_00795*△::*NAT* | 4 |
| *Cryptococcus var. grubii* (VNI) | CK4965 | *MAT*α *CNAG_04007*△::*NAT* | 4 |
| *Cryptococcus var. grubii* (VNI) | CK6149 | *MAT*α *CNAG_06312*△::*NAT* | 4 |
| *Cryptococcus var. grubii* (VNI) | CK0204 | *MAT*α *CNAG_03741*△::*NAT* | 4 |
| *Cryptococcus var. grubii* (VNI) | CK5853 | *MAT*α *CNAG_05729*△::*NAT* | 4 |
| *Cryptococcus var. grubii* (VNI) | CK5639 | *MAT*α *CNAG_05305*△::*NAT* | 4 |
| *Cryptococcus var. grubii* (VNI) | CK6557 | *MAT*α *CNAG_07399*△::*NAT* | 4 |
| *Cryptococcus var. grubii* (VNI) | CK1916 | *MAT*α *CNAG_01806*△::*NAT* | 4 |
| *Cryptococcus var. grubii* (VNI) | CK4861 | *MAT*α *CNAG_03833*△::*NAT* | 4 |
| *Cryptococcus var. grubii* (VNI) | CK4113 | *MAT*α *CNAG_02339*△::*NAT* | 4 |
| *Cryptococcus var. grubii* (VNI) | CK4200 | *MAT*α *CNAG_02499*△::*NAT* | 4 |
| *Cryptococcus var. grubii* (VNI) | CK4496 | *MAT*α *CNAG_03107*△::*NAT* | 4 |
| *Cryptococcus var. grubii* (VNI) | CK1262 | *MAT*α *CNAG_04861*△::*NAT* | 4 |
| *Cryptococcus var. grubii* (VNI) | CK1076 | *MAT*α *CNAG_01653*△::*NAT* | 4 |
| *Cryptococcus var. grubii* (VNI) | CK2618 | *MAT*α *CNAG_01816*△::*NAT* | 4 |
| *Cryptococcus var. grubii* (VNI) | CK0844 | *MAT*α *CNAG_06082*△::*NAT* | 4 |
| *Cryptococcus var. grubii* (VNI) | NRHc5010.ENRα | *MAT*α Clinical isolate | 21 |
| *Cryptococcus var. grubii* (VNI) | Bt99α | *MAT*α Clinical isolate | 21 |
| *Cryptococcus var. grubii* (VNI) | MW-RSA82α | *MAT*α Clinical isolate | 21 |
| *Cryptococcus var. grubii* (VNI) | Bt49α | *MAT*α Clinical isolate | 21 |
| *Cryptococcus var. grubii* (VNI) | C23α | *MAT*α Clinical isolate | 21 |
| *Cryptococcus var. grubii* (VNI) | Bt130**a** | *MAT***a** Clinical isolate | 21 |
| *Cryptococcus var. grubii* (VNI) | Bt11α | *MAT*α Clinical isolate | 21 |
| *Cryptococcus var. grubii* (VNI) | NRHc5001.ENRα | *MAT*α Clinical isolate | 21 |
| *Cryptococcus var. grubii* (VNII) | LP-RSA1684α | *MAT*α Clinical isolate | 21 |
| *Cryptococcus var. grubii* (VNII) | PMHc1023.ENRα | *MAT*α Clinical isolate | 21 |
| *Cryptococcus var. grubii* (VNBI) | PMHc1026.ENRα | *MAT*α Clinical isolate | 21 |

| | | | |
|---|---|---|---|
| *Cryptococcus var. grubii* (VNBI) | NRHc5005.ENRα | *MAT*α Clinical isolate | 21 |
| *Cryptococcus var. grubii* (VNBI) | Muc364-1α | *MAT*α Mopane tree | 21 |
| *Cryptococcus var. grubii* (VNBI) | LP-RSA6618α | *MAT*α Clinical isolate | 21 |
| *Cryptococcus var. grubii* (VNBI) | Gbc573-1α | *MAT*α Environmental isolate | 21 |
| *Cryptococcus var. grubii* (VNBI) | Gbc573-1α | *MAT*α Environmental isolate | 21 |
| *Cryptococcus var. grubii* (VNBI) | Ftc103-1α | *MAT*α Environmental isolate | 21 |
| *Cryptococcus var. grubii* (VNBI) | Bt7α | *MAT*α Clinical isolate | 21 |
| *Cryptococcus var. grubii* (VNBI) | PMHc1011.ENR**a** | *MAT***a** Clinical isolate | 21 |
| *Cryptococcus var. grubii* (VNBI) | NRHc5020.ENR**a** | *MAT***a** Clinical isolate | 21 |
| *Cryptococcus var. grubii* (VNBI) | NRHc5004.ENR**a** | *MAT***a** Clinical isolate | 21 |
| *Cryptococcus var. grubii* (VNBI) | Muc498-2**a** | *MAT***a** Environmental isolate | 21 |
| *Cryptococcus var. grubii* (VNBII) | PMHc1052.ENR.STOR**a** | *MAT***a** Clinical isolate | 21 |
| *Cryptococcus var. grubii* (VNBII) | LP-RSA3144α | *MAT*α Clinical isolate | 21 |
| *Cryptococcus var. grubii* (VNBII) | PMHc1002.ENRα | *MAT*α Clinical isolate | 21 |
| *Cryptococcus var. grubii* (VNBII) | Muc387-1α | *MAT*α Environmental isolate | 21 |
| *Cryptococcus var. grubii* (VNBII) | MW-RSA913α | *MAT*α Clinical isolate | 21 |
| *Cryptococcus var. grubii* (VNBII) | Bt103α | *MAT*α Clinical isolate | 21 |
| *Cryptococcus var. grubii* (VNBII) | Bt55α | *MAT*α Clinical isolate | 21 |
| *Cryptococcus var. grubii* (VNBII) | Bt1α | *MAT*α Clinical isolate | 21 |
| *Cryptococcus var. grubii* (VNBII) | PMHc1024.ENR**a** | *MAT***a** Clinical isolate | 21 |
| *Cryptococcus var. grubii* (VNBII) | NRHc5014.ENR**a** | *MAT***a** Clinical isolate | 21 |
| *Cryptococcus var. grubii* (VNBII) | Bt81**a** | *MAT***a** Clinical isolate | 21 |
| *Cryptococcus var. grubii* (VNBII) | Bt65**a** | *MAT***a** Clinical isolate | 21 |
| *Cryptococcus var. grubii* (VNBII) | Bt133**a** | *MAT***a** Clinical isolate | 21 |
| *Cryptococcus var. grubii* (VNBII) | Bt206**a** | *MAT***a** Clinical isolate | 21 |
| *Cryptococcus deneoformans* | JEC20 | *MAT***a** wild type | 22 |
| *Cryptococcus deneoformans* | JEC21 | *MAT*α wild type | 22 |
| *Cryptococcus deneoformans* | XL280α | *MAT*α wild type | 23 |
| *Cryptococcus deneoformans* | XL280a | *MAT***a** wild type | 23 |
| *Cryptococcus gattii* (VGI) | WM276 | *MAT*α wild type | 24 |
| *Cryptococcus gattii* (VGI) | E566 | *MAT***a** wild type | 25 |
| *Cryptococcus deuterogattii* (VGII) | R265a | *MAT***a** wild type | 26 |
| *Cryptococcus deuterogattii* (VGII) | R265α | *MAT*α wild type | 26 |
| *Cryptococcus deuterogattii* (VGII) | CBS10090 | *MAT*α wild type | 27 |
| *Cryptococcus bacillisporus* (VGIII) | CA1280 | *MAT*α wild type | 28 |
| *Cryptococcus bacillisporus* (VGIII) | CA1873 | *MAT***a** wild type | 28 |
| *Cryptococcus tetragattii* (VGIV) | IND107 | *MAT*α wild type | 29 |
| *Cryptococcus tetragattii* (VGIV) | CBS11718 | *MAT***a** wild type | 29 |

### Table S9. Oligonucleotides utilized for this study.

| Oligo Name | Sequence 5' to 3' | Purpose |
|---|---|---|
| JOHE56967/YES605 | GCACCATGCCCAGACCTCC | RT-PCR for *HXL1* |
| JOHE56968/YES606 | ACCGCTGCCTGCATCTCC | |
| JOHE56349/YES444 | TACGTCGCTCTCGACTTTGA | RT-PCR/qPCR for *ACT1* |
| JOHE56350/YES445 | CCAAGGAGAGAAGGCTGGAA | |
| JOHE56844/YES584 | CTCTGAGGACGACAAGGACA | qPCR for *KAR2* |
| JOHE56845/YES585 | AGCTCAGAAAGCTGCTCCTC | |
| JOHE56848/YES588 | GTCGCTATCCAGGGACATTT | qPCR for *ALG7* |
| JOHE56849/YES589 | GTCCAAAGAGTTGAGGGCAT | |
| JOHE56850/YES590 | GCGGAGCCAAACTCTTCTCAGCC | Spliced *HXL1* integration at the safe-heaven locus |
| JOHE56851/YES591 | ATAATTCCCCCTCGCCCGCTG CTGAGTGGCAAGCGACGGGG | |
| JOHE56852/YES592 | CCCCGTCGCTTGCCACTCAG CAGCGGGCGAGGGGGAATTAT | |
| JOHE56719/YES531 | GGTCCTTTCATCTGGCTCTTCGCC | |
| JOHE56811/YES577 | gcgtac GCGGCCGC ACCACCCGTGACTGTTGC | |
| JOHE56695/YES525 | aatt CTCGAG CATCCCTACGCGAGCCAAC | |
